# Dopamine transmission at D1 and D2 receptors in the nucleus accumbens contributes to the expression of incubation of cocaine craving

**DOI:** 10.1101/2024.06.26.600812

**Authors:** Sophia J. Weber, Alex B. Kawa, Alana L. Moutier, Madelyn M. Beutler, Lara M. Koyshman, Cloe D. Moreno, Jonathan G. Westlake, Amanda M. Wunsch, Marina E. Wolf

**Affiliations:** Department of Behavioral Neuroscience, Oregon Health & Science University, Portland, OR 97239

## Abstract

Relapse represents a consistent clinical problem for individuals with substance use disorder. In the incubation of craving model of persistent craving and relapse, cue-induced drug seeking progressively intensifies or ‘incubates’ during the first weeks of abstinence from drug self-administration and then remains high for months. Previously, we and others have demonstrated that expression of incubated cocaine craving requires strengthening of excitatory synaptic transmission in the nucleus accumbens core (NAcc). However, despite the importance of dopaminergic signaling in the NAcc for motivated behavior, little is known about the role that dopamine (DA) plays in the incubation of cocaine craving. Here we used fiber photometry to measure DA transients in the NAcc of male and female rats during cue-induced seeking tests conducted in early abstinence from cocaine self-administration, prior to incubation, and late abstinence, after incubation of craving has plateaued. We observed DA transients time-locked to cue-induced responding but their magnitude did not differ significantly when measured during early versus late abstinence seeking tests. Next, we tested for a functional role of these DA transients by injecting DA receptor antagonists into the NAcc just before the cue-induced seeking test. Blockade of either D1 or D2 DA receptors reduced cue-induced cocaine seeking after but not before incubation. We found no main effect of sex in our experiments. These results suggest that DA contributes to incubated cocaine seeking but the emergence of this role reflects changes in postsynaptic responsiveness to DA rather than presynaptic alterations.

## INTRODUCTION

A major difficulty in treating substance use disorder (SUD) is the propensity for relapse in abstinent individuals. ‘Incubation of craving’ is a leading preclinical model for studying vulnerability to cue-induced relapse after protracted abstinence from drug self-administration. It is based on the discovery that cue-induced cocaine seeking in rats progressively increased (“incubated”) over the first month or so of abstinence and then remained high for additional months before declining, providing a model for persistent vulnerability to relapse [1-3]. Incubation has subsequently been demonstrated for other addictive drugs [4-6] and in humans [7-10].

Although many brain regions contribute to incubation of cocaine craving [6,11], our lab and others have shown that synaptic plasticity leading to strengthening of excitatory synapses on medium spiny neurons (MSN) in the NAc core (NAcc) and shell (NAcs) subregions is necessary for its expression [11,12]. In contrast, very little is known about the role of NAc dopamine (DA) in incubation of cocaine craving. This is a significant knowledge gap since DA transmission in the NAc is essential for stimulus-reward learning and motivated behavior [13-16]. Most relevant to the incubation model, microdialysis and voltammetry studies have shown that DA release is associated with presentations of reward-related cues and the ability of such cues to promote reward seeking, both for natural rewards [e.g., 17,18,19] and cocaine [e.g., 20,21,22]. More recently, fiber photometry studies have detected DA transients in NAcc during cue-induced reinstatement of cocaine seeking [23] and in lateral shell in association with context-induced cocaine seeking [24].

Surprisingly, no studies have measured DA release during expression of cocaine incubation. Here we used fiber photometry paired with the DA biosensor GRAB_DA2m [25,26] to assess DA release in the NAcc during cue-induced seeking tests following long-access cocaine self-administration on forced abstinence day (FAD) 1-2, prior to incubation, and FAD40-50, after incubation. Then we tested the functional significance of this DA release by infusing DA D1-class and D2-class receptor (D1R and D2R) antagonists into NAcc prior to seeking tests.

## MATERIALS AND METHODS

Details are in Supplementary Methods.

### Subjects and Surgery

Procedures followed the NIH Guide for the Care and Use of Laboratory Animals and were approved by the OHSU IACUC. Male and female Long-Evans rats were used (Table S1). Rats for fiber photometry received jugular catheters, bilateral infusions of AAV9-hSyn-GRAB_DA2m or AAV9-hSyn-GRAB_DAmut into NAcc, and fiber optic cannula implantation above each infusion site. Rats for pharmacology experiments were implanted with jugular catheters and bilateral 23-gauge guide cannula above the NAcc. After recovery, rats underwent cocaine self-administration training (0.5 mg/kg/infusion; 6 h/day x 10 days).

### Fiber photometry recordings

During cue-induced seeking tests on FAD1-2 or FAD40-50, we used the TDT RZ10x system to record DA and isosbestic channels.

### Intra-NAcc microinjections

Rats received the D1R antagonist SCH39166, the D2R antagonist L-741,626, or vehicle prior to cue-induced seeking or open field tests.

### Analysis

Fiber photometry data were analyzed using GuPPy [27]. Full statistical analysis of processed photometry data and behavioral data are shown in Supplemental Tables 2-10.

## RESULTS

### Expression of GRAB_DA2m does not alter cocaine self-administration or incubation of craving

As our experiments are long in duration (see Timeline in Fig. 1A), we began by determining the stability of virus expression. Drug-naïve rats received NAcc infusion of an AAV expressing GRAB_DA2m and were killed after 3 or 10 weeks to parallel timing of cue-induced seeking tests in subsequent experiments. Immunohistochemical analysis demonstrated stable expression over this period (Fig. 1B; *t*_*11*_=0.110, *p*=0.915; Table S2 contains this and all other statistical analyses for Fig. 1).

**Figure 1.**
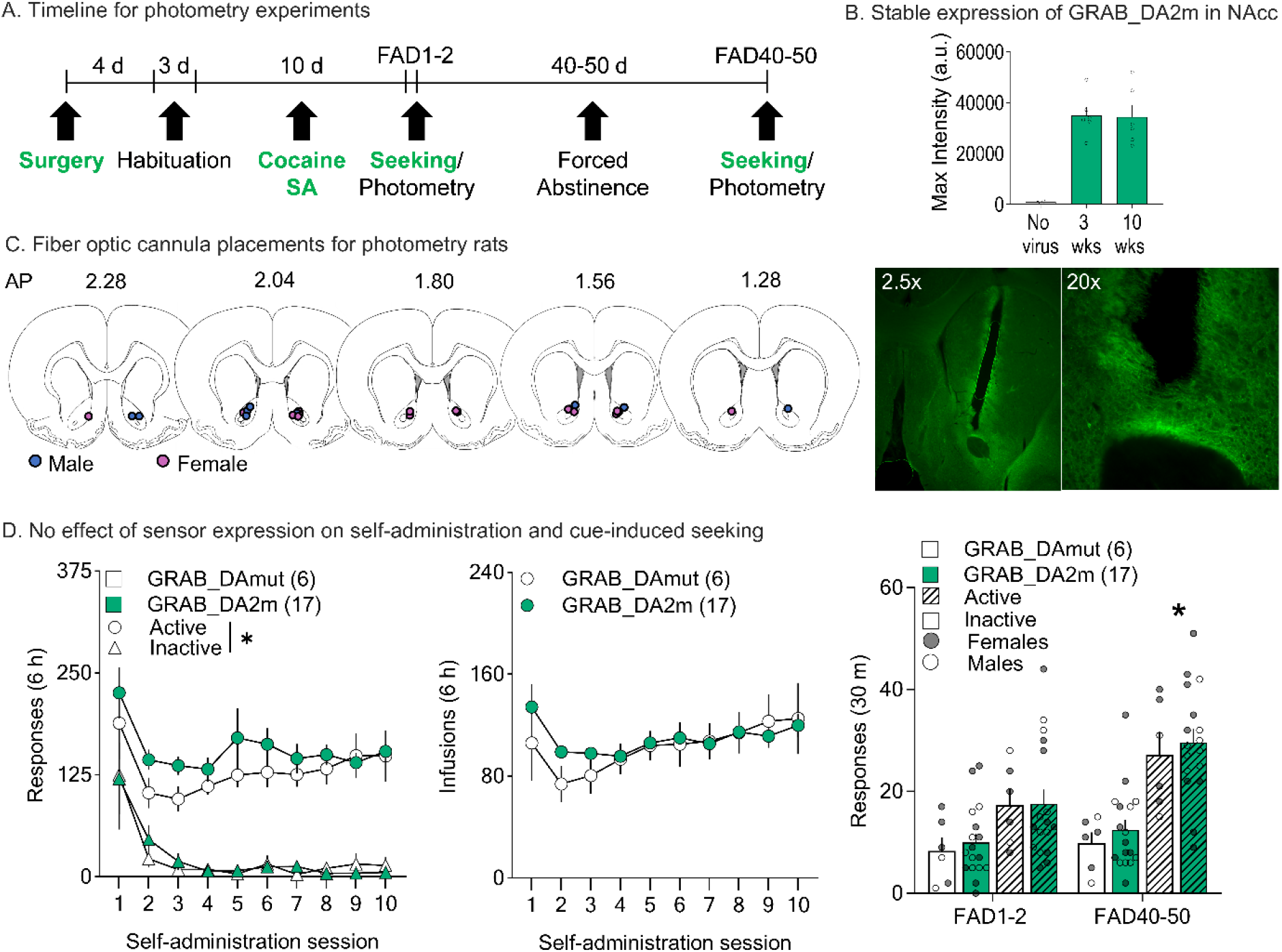
Behavior and virus expression in rats destined for fiber photometry. **A**. Timeline for photometry experiment (green denotes stages shown in this figure). **B**. *Top:* AAV9-hSyn-GRAB_DA2m expression in NAcc quantified (mean ± SEM) over time in three groups: no virus control group (n = 2 hemispheres), 500nL of virus assessed after 3 weeks of expression (n = 3 rats/6 hemispheres), and 500nL of virus after 10 weeks of expression (n = 3 rats/6 hemispheres). Dots show values for individual hemispheres. *Bottom:* Representative image after 3 weeks of expression at two different magnifications. **C**. Fiber optic cannula placement for photometry rats. The dot signifies the end of the cannula (females, pink; males, blue). Placement was determined after immunohistochemistry for GFP to confirm virus expression at the end of the cannula. Sections adapted from Paxinos & Watson 7^th^ edition. **D**. Cocaine self-administration and cue-induced seeking data. *Left:* Rats expressing either GRAB_DA2m (n = 17) or GRAB_DAmut (n = 6) learned to nose poke into the active port for an infusion of cocaine and discriminated between the drug-associated (active) port and the inactive port (*p<0.05; Table S2). *Middle:* Cocaine infusions for sensor and mutant expressing rats during 10 days of self-administration training. Mean (±SEM) for each 6-h session is shown. *Right:* Cue-induced seeking tests on FAD1-2 and FAD40-50 for sensor and mutant expressing rats. Bars show mean (±SEM) for each 30-min seeking test while dots indicate individual rats (open circles, males; closed circles, females). * indicates significant main effect of test day but not virus for active pokes on FAD40-50 versus FAD1-2 (Table S2). AP, anterior posterior; FAD, forced abstinence day.

Next, we tested whether expression of the sensor, which potentially could lead to buffering of DA levels, might be affecting our measured behaviors. Rats underwent intra-NAcc infusion of GRAB_DA2m or a mutant version that does not bind DA (GRAB_DAmut), implantation of a fiber optic cannula above the area of virus expression, and implantation of a jugular catheter, followed by cocaine self-administration training (Fig. 1A, C). During training, nose-pokes in the active port triggered cocaine infusion, a 4-sec light cue/simultaneous 4-sec time-out period. Both virus groups acquired cocaine self-administration and earned a similar number of infusions (Fig. 1D). Analysis revealed significant effects of session and port, but not virus. For one rat, inactive port data on days 6-7 were excluded as outliers; see Fig. S1 for full data. After 10 days of self-administration, rats received cue-induced seeking tests on FAD1-2 and FAD40-50 (within-subject design), during which pokes in the active port triggered the cue but no cocaine infusion (Fig. 1D-right). Mixed effects analysis revealed no effect of virus on drug seeking but an interaction between port and test day (*F*_1,21_=9.071, *p*=0.00664), indicative of incubation. Finally, we found no main effect of sex for any measure related to cocaine self-administration or cue-induced seeking (Fig. S2A).

### Responding for cocaine cues elicits similar DA responses in early and late abstinence

Fiber photometry recordings were performed in both GRAB_DA2m and GRAB_DAmut rats during FAD1-2 and FAD40-50 seeking tests (Fig. 2A). As expected, there was no apparent signal for GRAB_DAmut-expressing rats (Fig. S3), so we used their data to define analysis thresholds (see below).

**Figure 2.**
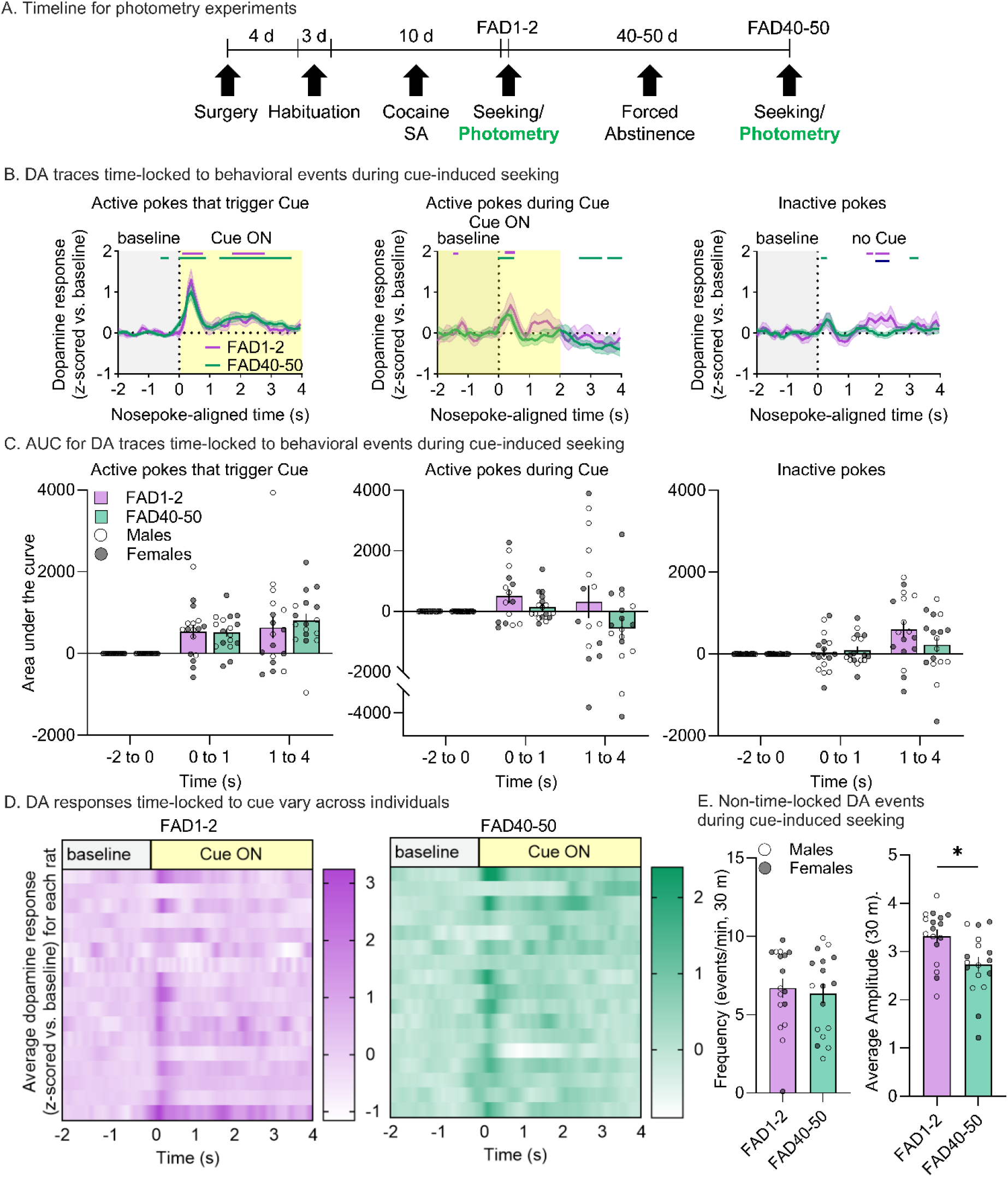
Fiber photometry recordings during cue-induced seeking tests. **A**. Timeline for photometry experiment (green denotes stages shown in this figure). **B**. DA traces time-locked to behavioral events (n = 17 rats). *Left:* z-scored mean DA response normalized to a baseline period (-2 to 0 s) for active pokes that triggered the cue on FAD1-2 and FAD40-50. SEM is shown in shaded area around the mean. The matching-colored lines above the traces show periods in the 6-sec window during which bootstrapping indicates 95% confidence that the mean is not equal to zero. *Middle:* z-scored DA response for active pokes during cue light on/time-out period on FAD1-2 and FAD40-50. *Right:* z-scored DA response for inactive pokes on FAD1-2 and FAD40-50. **C**. Area under the curve for traces shown in (B). Traces are split into time epochs of baseline (-2 to 0 s), initial peak (0 to 1 s) and secondary peak (1 to 4 s). Bars show mean (±SEM) for each time period around the behavioral event while dots indicate individual rats. **D**. Heat maps showing the average DA trace for each rat that contributed to (B). *Left:* DA responses to active pokes that trigger the cue for individual rats on FAD1-2. *Right:* Individual responses on FAD40-50. Darker colors represent higher z-score. **E**. Analysis of DA events not time-locked to a behavioral event. Data obtained using the mutant sensor GRAB_DA2mut were used to define a threshold for DA transient identification such that 0 peaks were detected in the mutant trace; this threshold was applied to the entire 30 min recording for each seeking test for all rats (n = 17). *Left:* Frequency (average number of events per minute) on FAD1-2 and FAD40-50. *Right:* Average amplitude of all events above threshold in the 30 min recording for FAD1-2 and FAD40-50. Bars show mean (±SEM) for each 30-min seeking test while dots indicate individual rats (open circles, males; closed circles, females). *p<0.05 FAD1-2 versus FAD40-50 (Table S4). AUC, area under the curve; FAD, forced abstinence day.

Photometry data for GRAB_DA2m-expressing rats on FAD1-2 and FAD40-50 are presented in Fig. 2B. Full statistical analysis for these and other data in Fig. 2 are presented in Tables S4 and S6. DA levels, plotted relative to each nose-poke (time 0), are shown for pokes in the active port that triggered the cue light (Fig. 2B-left), pokes in the active hole that did not trigger cue presentation because they occurred during the 4-s time-out period when the cue light is already on (Fig. 2B-center), and pokes in the inactive port (Fig. 2B-right). Horizontal lines above the traces indicate DA response duration assessed using continuous threshold bootstrapping methods, where the 95% confidence interval does not cross baseline (z-score=0) for the consecutive threshold period [28-30], with the consecutive threshold defined using GRAB_DAmut data (see Supplementary Methods). Periods during which bootstrapping revealed between-group differences are indicated by a dark blue line.

Focusing on active pokes triggering the cue, analysis of both FAD1-2 and FAD40-50 revealed an increase in DA levels at cue onset (0-1 s) as well as a smaller amplitude but longer-lasting increase during the remainder of cue light presentation (1-4 s) (Fig. 2B-left). Although there were small differences in the exact timing of DA increases (see colored bars above traces), we found no significant differences between test days when comparing bootstrapped confidence intervals. We also quantified both periods relative to baseline (-2 to 0 s) via area under the curve (AUC) (Fig. 2C-left). This revealed a significant effect of time (*F*_2,32_=14.0, *p*<0.0001) but not test day (FAD1-2 vs FAD40-50) and no significant interaction. These data indicate that, compared to baseline, active pokes triggering the cue elicit DA release that is of similar magnitude on FAD1-2 and FAD40-50 when averaged across trials and subjects. However, heat maps showing the average DA response for each subject revealed considerable variability (Fig. 2D). This variability could have may origins, including individual differences in drug responding [e.g., 31] and the area of NAcc targeted [32].

Next, we performed the same analyses for active port nose-pokes made within the time-out period (Cue light on) (Fig. 2B-middle, Tables S4 and S6). The pattern of DA responses was similar to that in Fig. 2B-left, but less robust, with only the initial response detected by bootstrapping. Again, no difference between test days was indicated. Mixed effects analysis of AUC for 0-1 s and 1-4 s time periods (relative to -2 to 0 s baseline) found no significant effect of time or test day (Fig. 2C-middle).

We also examined the DA response to inactive pokes (Fig. 2B-right, Tables S4 and S6). Here, bootstrapping identified a very brief increase shortly after the nose-poke on FAD40-50, and a more delayed response (∼ 2 sec) that differed significantly between test days. Analysis of AUC revealed a significant effect of time (*F*_2,32_=6.97, *p*=0.00307) but not test day and no significant interaction (Fig. 2C-right). Overall, Fig. 2B suggests that the DA response immediately associated with the nose-poke (0-1 s) is greater for active than inactive pokes.

We analyzed frequency and amplitude of all DA transients over the 30-min test, regardless of whether they were time-locked to a nose-poke (Fig. 2E, Table S4). We used GRAB_DA2mut data to define a threshold for DA transient identification such that 0 peaks were detected in the mutant trace after applying this threshold (Supplementary Methods). The frequency of DA transients did not differ between test days, but the average amplitude of all above-noise DA events was significantly reduced on FAD40-50 versus FAD1-2 (*t*_16_=2.84, *p*=0.0117). This could reflect alterations in tonic DA release as a function of abstinence time and/or a response to the context previously paired with cocaine self-administration.

We investigated within-session effects by binning seeking test data into three 10-min bins. Behaviorally, we observed greater active port responses across bins on FAD40-50 vs FAD1-2, reflecting incubation of craving (Fig. 3B, Table S2; main effect of test day: *F*_1,17_=10.7, *p*=0.0001). For photometry data, variability was observed across bins on FAD1-2, with the highest magnitude DA response in the first 10 min, whereas responses in each bin were similar on FAD40-50 (Fig. 3C). A concern when comparing the two test days is that we averaged different numbers of DA responses. Thus, because of the incubation phenomenon, rats made more active port responses on FAD40-50 than FAD1-2. This would be expected to reduce variability for the FAD40-50 test. To control for this, we analyzed the DA response to the first nose-poke in each 10-min bin, so that each rat has only one poke/DA response contributing to each bin for each test day. When analyzed in this manner, the pattern of responding across bins was similar on FAD1-2 and FAD40-50 (Fig. 3D). We also compared the DA response to the first nose-poke of each session and observed no difference between test days (Fig. 3E; Table S5). Thus, within-session DA responses do not differ between test days when accounting for the number of responses in the data-sets.

**Figure 3.**
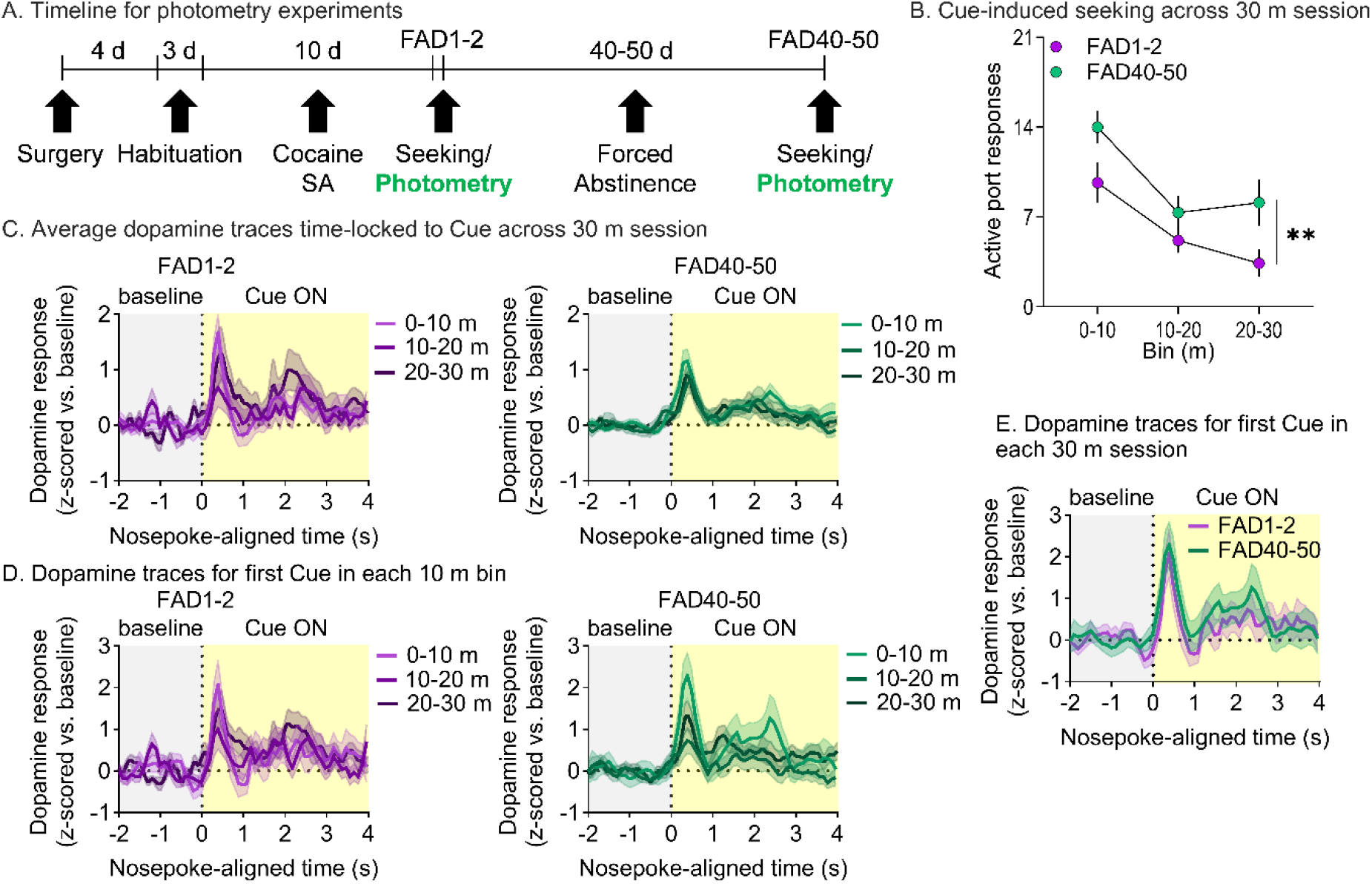
Analysis of behavior and DA responses during specific periods of cue-induced seeking tests. **A**. Timeline for photometry experiment (green denotes stages shown in this figure). **B**. Mean (±SEM) active port responses for seeking test data separated into 10-min bins (n = 17 rats in this and all subsequent panels). **C**. Mean DA trace (±SEM) time-locked to active pokes that triggered the cue, split into 10-min bins, on FAD1-2 (*Left*) and FAD40-50 (*Right*). Separate bins are shown as different colored lines. **D**. Mean DA trace (±SEM) time-locked to the first poke that triggered a cue in each 10-min bin shown in (C). Separate bins are shown as different colored lines. **E**. Mean DA trace (±SEM) time-locked to the first poke that triggered a cue in each 30-min session for FAD1-2 and FAD40-50. FAD, forced abstinence day.

Males and females exhibited a similar pattern of DA release during seeking tests (no main effect of sex: *F*_1,30_=1.10, *p*=0.304), although small time-dependent differences between the sexes were suggested by disaggregated data (Fig. S2B, Table S5). Although there is evidence that DA transmission is modulated by the estrous cycle [33,34], very few females were in estrus at the time of testing; therefore, we cannot make conclusions about the influence of estrous cycle (Fig. S2C).

### Intra-NAcc administration of a D1R antagonist decreases expression of cocaine incubation

To determine whether DA responses during cue-induced seeking tests have behavioral significance, we infused D1-or D2-receptor antagonists into NAcc 15 min prior to cue-induced seeking tests on FAD1 or FAD40-50. Results obtained with the D1 antagonist SCH39166 are shown in Fig. 4 (full statistics in Table S7). After self-administration training, rats were divided into equivalent groups, destined for vehicle or SCH39166 infusion, based on cocaine infusions on the last three self-administration days (sessions 8-10) (Fig. 4B-left). For rats assigned to the FAD40-50 test group, vehicle and SCH39166 groups were also balanced based on results of the FAD1 seeking test (Fig. S4A).

**Figure 4.**
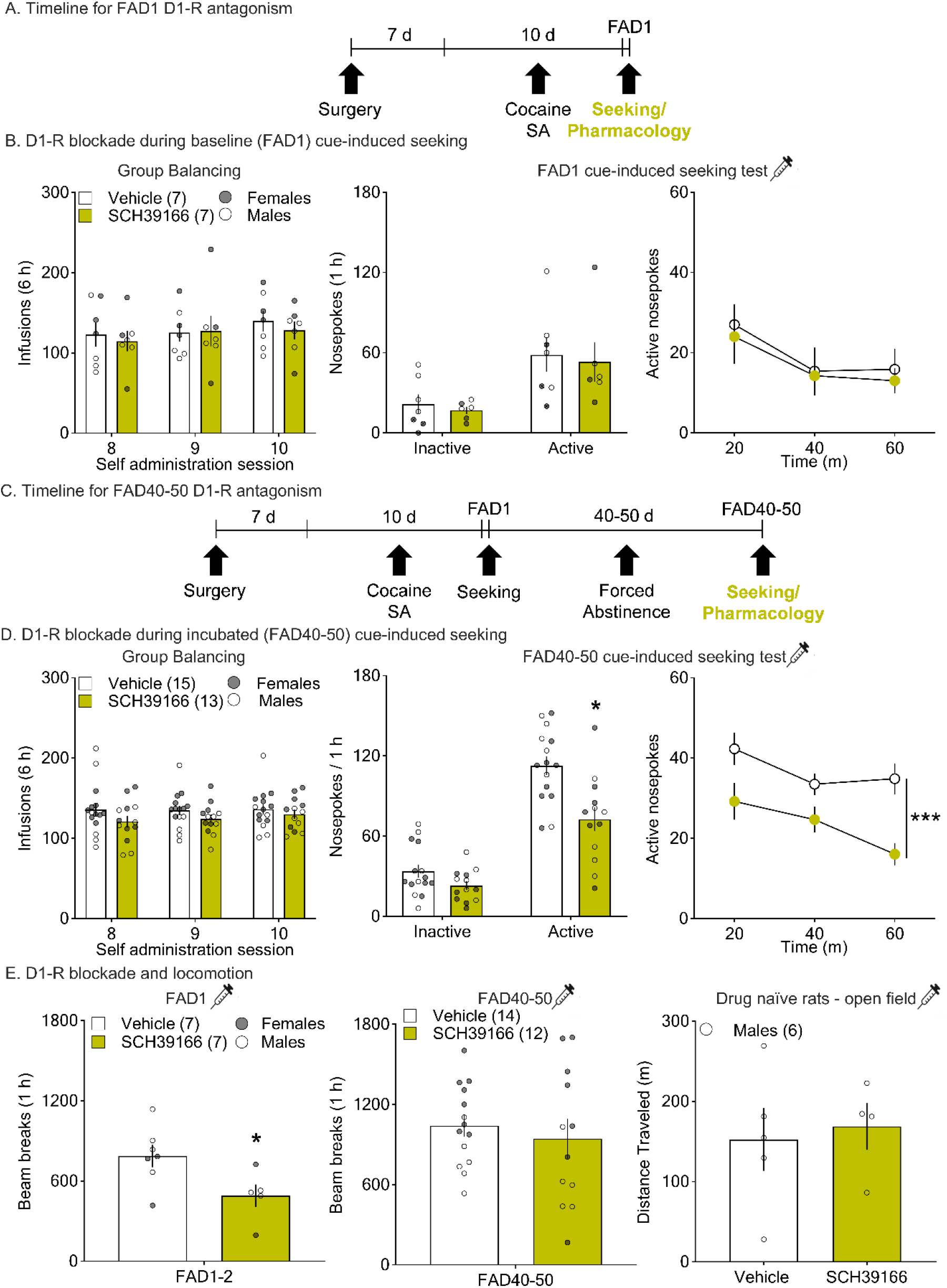
Intra-NAcc infusion of the D1R antagonist SCH39166 reduces cue-induced seeking on FAD40-50 but not FAD1. **A**. Timeline for FAD1 D1R antagonist experiment (yellow denotes stages that are the focus panel B). **B**. Behavioral data for FAD1 D1R antagonist experiment. *Left:* Groups destined for vehicle or SCH39166 infusion on FAD1 were balanced based on mean (±SEM) cocaine infusions for the last three 6-h sessions of self-administration training. *Middle:* Bars show mean (±SEM) active and inactive pokes during the 1-h FAD1 cue-induced seeking test for rats receiving intra-NAcc vehicle (n = 7) or SCH39166 (n = 7). Dots indicate individual rats (open circles, males; closed circles, females). *Right:* Mean (±SEM) active pokes for FAD1 seeking test data split into three 20-min bins. **C**. Timeline for FAD40-50 D1R antagonist experiment (yellow denotes stages that are the focus of panel D). **D**. Behavioral data for FAD40-50 D1R antagonist experiment. *Left:* Groups destined for vehicle or SCH39166 infusion on FAD40-50 were balanced based on mean (±SEM) cocaine infusions for the last three 6-h sessions of self-administration training (shown here) as well as FAD1 seeking data (Fig. S4A). *Middle:* Mean (±SEM) pokes during the 1-h FAD40-50 cue-induced seeking test for rats receiving intra-NAcc vehicle (n = 15) or SCH39166 (n = 13) (*p<0.05 vs vehicle; Table S7). *Right:* Mean (±SEM) active pokes for FAD40-50 seeking test data split into three 20-min bins. Incubated cue-induced seeking was suppressed in the SCH39166 group (***p<0.001 main effect of treatment, SCH39166 vs vehicle; Table S7). **E**. Operant box photobeam breaks during seeking tests and a separate open field experiment conducted in drug-naïve rats. *Left:* Mean (±SEM) beam breaks during the FAD1 seeking test shown in (B) (n = 7 rats/group). *Middle:* Mean (±SEM) beam breaks during the FAD40-50 seeking test shown in (D) (n = 14 for vehicle, n = 12 for SCH39166; two operant boxes lacked functional photobeams). *Right:* Mean (±SEM) total distance traveled in meters (m) during a 1-h open field test conducted after intra-NAcc injection of vehicle or SCH39166 in drug-naïve rats (n = 6 total; 3 rats received both vehicle and SCH39166 in a counter-balanced design with one day between tests, 2 rats received only vehicle, and 1 rat received only SCH39166). SCH39166 significantly reduced beam breaks given prior to the FAD1 test (*p<0.05 vs vehicle), but not when given before the FAD40-50 test and had no effect on open field locomotion (Table S7). Dots show data for individual rats. FAD, forced abstinence day.

Infusion of SCH39166 (1 μg/0.5 μL) in NAcc (placements shown in Fig. S5) did not affect cue-induced seeking on FAD1 compared to vehicle-infused rats (Fig. 4B-right). When we binned the 1-h test into three 20-min bins, a mixed effects analysis showed a significant effect of bin (*F*_2,24_=5.21, *p*=0.0132) but not treatment and no significant treatment x bin interaction. In contrast, when the D1R antagonist was infused prior to a FAD40-50 test, it significantly suppressed incubated cue-induced seeking (Fig. 4D-middle; treatment x port: *F*_1,26_=6.03, *p*=0.0210) specifically on the active port (*t*_52_=4.44, p<0.0001). Analysis of binned data from FAD40-50 (Fig. 4D-right) revealed a significant effect of treatment (*F*_1,26_=16.7, *p*=0.0124) and bin (*F*_2,52_=4.79, *p*=0.000377), but no interaction. We found no main effect of sex when analyzing aggregated data from this experiment (Table S8), although disaggregated data may suggest a more robust effect in males (Fig. S4B). Overall, these data indicate that D1R blockade significantly reduced seeking on FAD40-50 but not FAD1.

Because DA receptor antagonists could reduce drug seeking via a general reduction in motor behavior, we monitored locomotor activity during seeking tests using photobeams in operant chambers. SCH39166 significantly reduced beam-breaks on FAD1 (Fig. 4E-left; *t*_10_=2.43, p=0.0352) but not FAD40-50 (Fig. 4E-middle). We also tested the effect of intra-NAcc vehicle or SCH39166 (same dose) on open field behavior in drug-naïve rats and found no effect of treatment (Fig. 4E-right). These results, combined with no significant effect of SCH39166 on inactive nose-pokes (Fig. 4D-middle), suggest that the observed effect of D1R antagonism on FAD40-50 cue-induced seeking does not reflect a general reduction in motor behavior.

### Intra-NAc administration of a D2 antagonist decreases expression of cocaine incubation

We performed a parallel experiment using the highly specific D2R antagonist L-741,626 and its vehicle (Fig. 5; full statistics in Table S9). Intra-NAcc infusion of L-741,626 (1 μg/0.5 μL; placements shown in Fig. S7) did not affect cue-induced seeking when given prior to the FAD1 test (Fig. 5B-middle). There was also no effect of treatment when we split the 1-h FAD1 seeking test into 20-min bins (Fig. 5B-right). However, similar to D1R antagonism, we observed a significant suppression of incubated cue-induced seeking when L-741,626 was given on FAD40-50 (Fig. 5D-right; treatment x port interaction: *F*_1,21_=7.06, *p*=0.0148). Post hoc analysis confirmed that L-741,626 specifically reduced active port responding (*t*_42_=3.46, p=0.00251). We performed the same analysis after binning the data and found a main effect of treatment (*F*_1,21_=6.32, *p*=0.0202) and bin (*F*_2,42_=5.41, *p*=0.00811), but no interaction. Analysis of aggregated data from this experiment found no main effect of sex (Table S10), but disaggregated data may suggest a more robust effect of L-741,626 in males (Fig. S6B, Table S10). Overall, these data indicate that D2R blockade significantly reduced incubated cue-induced seeking on FAD40-50 but not FAD1 seeking.

**Figure 5.**
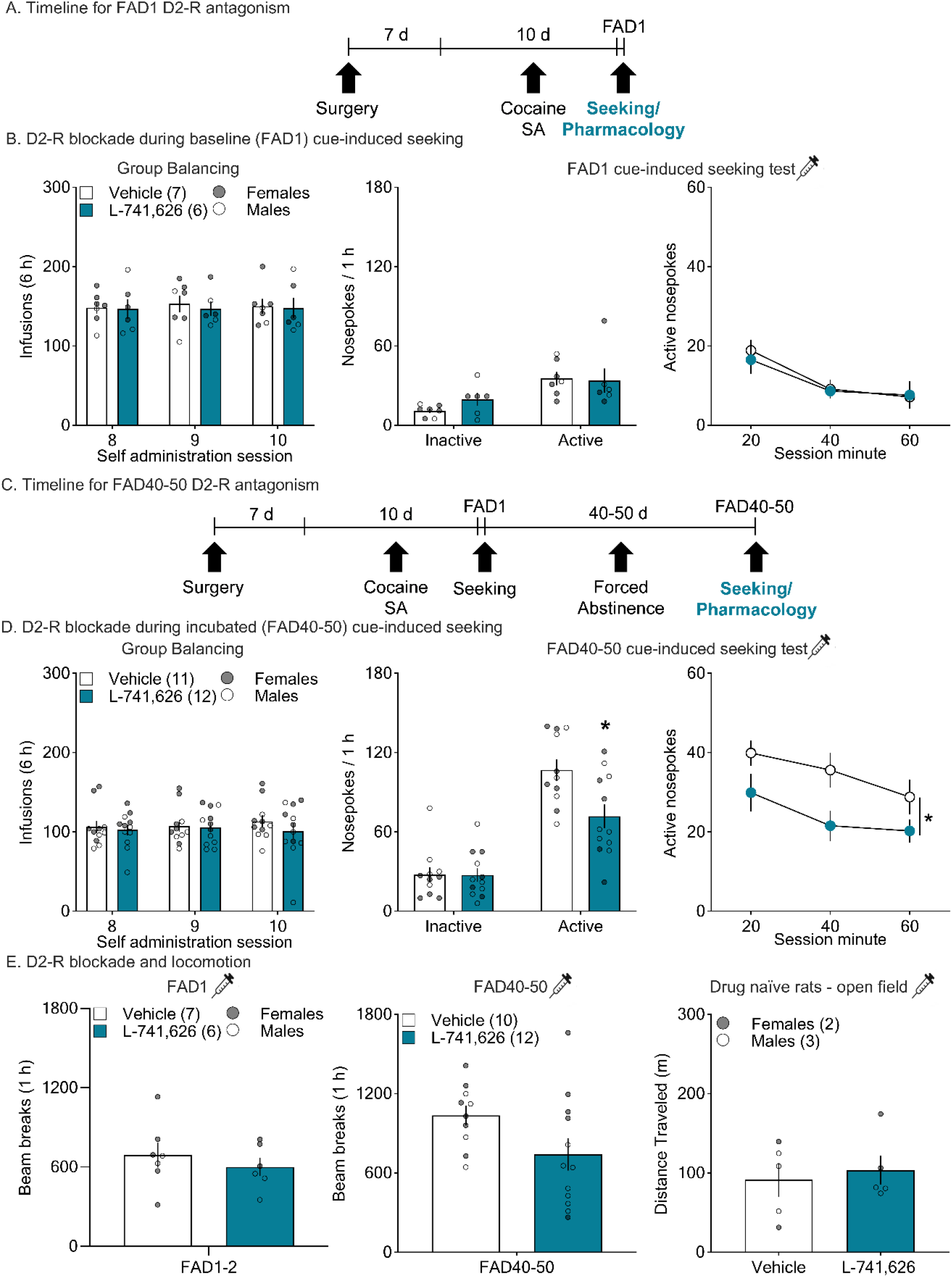
Intra-NAcc infusion of the D2R antagonist L-741,626 reduces cue-induced seeking on FAD40-50 but not FAD1. **A**. Timeline for FAD1 D2R antagonist experiment (teal denotes stages that are the focus of panel B). **B**. Behavioral data for FAD1 D2R antagonist experiment. *Left:* Groups destined for vehicle or L-741,626 infusion on FAD1 were balanced based on mean (±SEM) cocaine infusions for the last three 6-h sessions of self-administration training. *Middle:* Mean (±SEM) pokes for the 1-h FAD1 cue-induced seeking test for rats receiving intra-NAcc vehicle (n = 7) or L-741,626 (n = 6). Dots indicate individual rats (open circles, males; closed circles, females). *Right:* Mean (±SEM) active pokes for FAD1 seeking test data split into three 20-min bins. **C**. Timeline for FAD40-50 D2R antagonist experiment (teal denotes stages that are the focus of panel D). **D**. Behavioral data for FAD40-50 D2R antagonist experiment. *Left:* Groups destined for vehicle or L-741,626 infusion on FAD40-50 were balanced based on mean (±SEM) cocaine infusions for the last three 6-h sessions of self-administration training (shown here) as well as FAD1 seeking data (Fig. S6A). *Middle*: Mean (±SEM) pokes for the 1-h FAD40-50 cue-induced seeking test for rats receiving intra-NAcc vehicle (n = 11) or L-741,626 (n = 12) (*p<0.05 vs vehicle; Table S9). *Right:* Mean (±SEM) active pokes for FAD40-50 seeking test data split into three 20-min bins (*p<0.05 main effect of treatment, L-741,626 versus vehicle; Table S9). **E**. Operant box photobeam breaks during seeking tests and a separate open field experiment conducted in drug-naïve rats. *Left:* Mean (±SEM) beam breaks during the FAD1 seeking test shown in (B) (n = 7 for vehicle, n = 6 for L-741,626). *Middle:* Mean (±SEM) beam breaks during the FAD40-50 seeking test shown in (D) (n = 10 vehicle, n = 12 L-741,626; two operant boxes lacked functional photobeams). *Right:* Mean (±SEM) total distance traveled in meters (m) during a 1-h open field test conducted after intra-NAcc injection of vehicle or L-741,626 in drug-naïve rats (n = 5 total, all rats received both vehicle and L-741,626 in a counter-balanced design with one day between tests). FAD, forced abstinence day.

To test for effects of the D2R antagonist on locomotion, we analyzed beam breaks in the operant chamber during seeking tests and found no significant difference between the L-741,626 and vehicle groups on FAD1 or FAD40-50 (Fig. 5E-left and middle). We also tested a separate cohort of drug-naïve rats for open field locomotion and found no effect of treatment (Fig. 5E-right). These data, along with no effect of L-741,626 on inactive nose-pokes (Fig. 5D-middle), indicate that the effect of D2R antagonism on incubated cue-induced seeking is not due to motor impairment.

## DISCUSSION

During cue-induced seeking tests conducted during forced abstinence from cocaine self-administration, we used fiber photometry to detect DA transients time-locked to nose-pokes that triggered presentation of the cue previously paired with cocaine infusions. We found that the magnitude of this DA response did not differ significantly on FAD1-2 and FAD40-50. To test the functional significance of this DA release, we performed intra-NAcc injections of the D1R antagonist SCH39166 prior to FAD1 or FAD40-50 seeking tests. When given on FAD1, we found no effect of SCH39166 on active port responses. However, when SCH39166 was given on FAD40-50, after cocaine cue-induced seeking has incubated, we found a significant suppression of active port responses. We performed a parallel study with L-741,626, a specific D2R antagonist, and similarly found a selective suppression of incubated cue-induced seeking. Our results show for the first time that DA release occurs in the NAcc in conjunction with incubated cue-induced cocaine seeking and is required for its expression.

### The magnitude of cue-induced DA release is not altered over incubation although the significance of DA receptor stimulation for cue-induced seeking is changed

When measuring DA release during cue-induced cocaine seeking tests, and time-locking to the seeking response (active pokes that trigger the cue), we found that the average magnitude of DA release was the same on FAD1-2 and FAD40-50. This suggests that incubated cue-induced seeking is not related to an increase in DA release. However, given that DA-R blockade in the NAcc inhibits incubated seeking (FAD40-50) but not FAD1 seeking, we suggest that postsynaptic adaptations occur during incubation such that the same amount of DA release evokes a greater postsynaptic response. Supporting this possibility, it has long been known that adaptations in DA receptor expression and post-receptor intracellular signaling occur after cocaine exposure [35,36]. We previously examined surface expression of DA-Rs in NAcc after the same cocaine self-administration regimen used here and found no change in D1Rs and small decreases in D2Rs during cocaine incubation [37], so a change in post-receptor signaling seems more likely to account for our results.

We acknowledge that the relationship between the magnitudes of DA release and cue responding may depend on the design of the study. A recent study measured DA release in response to non-contingent presentation of a cue previously paired with cocaine self-administration and found greater DA release after 30 days of forced abstinence [38]. The difference likely reflects the fact that this study measured the DA response to non-contingent cue presentation whereas we measured the DA response to the seeking action paired with the cue. Furthermore, this study did not demonstrate a causal relationship between greater DA release in response to non-contingent cue presentation and the expression of incubated cocaine seeking. Another recent study measured DA release in the lateral NAc shell during cocaine seeking in mice and found an increase in the frequency of DA transients measured on FAD24 compared to pre-SA baseline [24]. We observed a change in amplitude rather than frequency, perhaps because our recordings were performed in NAcc or because we compared different time-points. Finally, in slice voltammetry studies performed in NAcc after 4 weeks of abstinence from cocaine SA regimens leading to incubation of craving, electrically stimulated DA release did not differ significantly from that observed in cocaine-naïve rats, although cocaine was more potent at DA uptake inhibition [39].

Some theories emphasize the significance of DA release not only for learning about cues and rewards but also for encoding motivational vigor and saliency [40-43]. Our data are generally consistent with these findings although our behavioral task is not designed to isolate these facets of DA-related behavior.

### Both D1 and D2 receptors contribute to incubated cue-induced cocaine seeking

Our studies revealed that blocking either the D1R or D2R similarly attenuated incubated cue-induced cocaine seeking. These results are consistent with recent models supporting cooperative interactions between D1R expressing MSN (D1-MSN) and D2R expressing MSN (D2-MSN) in regulating motivated behavior [44-46]. It is also important to keep in mind that our pharmacology experiments were focused at the level of D1R and D2R transmission, not D1-MSN and D2-MSN activity. We also acknowledge that DA receptors are present not only on MSN (where they act as neuromodulators to influence excitability, glutamate transmission, and intracellular signaling) but also on other neuronal elements, indicating a multiplicity of potential interactive regulatory mechanisms [47].

While our study is the first to show that activation of D1R and D2R is required for cocaine incubation, a previous study found that activation of both receptors in NAcc but not shell is required for expression of incubation of methamphetamine craving [48]. Others found that activation of both D1R and D2R in NAc shell is necessary and sufficient for drug-primed reinstatement of cocaine seeking [49,50]. Overall, these results support cooperative actions of DA signaling at D1R and D2R in psychostimulant seeking. Consistent with these findings at the level of the NAc, functional inhibition of VTA DA neurons reduces cocaine reinstatement [51] and cocaine seeking after brief abstinence [52].

While we focused on D1R and D2R as these are the most widely expressed DA receptors in NAcc, some NAc MSN also express the D3R [53]. Indeed, one study found that a D3R antagonist reduced context-induced cocaine seeking during forced abstinence, although this occurred independent of abstinence duration [54]. Based on this study, it is unlikely that D3R contribute to observed effects of the D2R antagonist L-741,626 on cocaine seeking, as our effects were specific to incubated cue-induced cocaine seeking. However, we acknowledge that there can be differences in mechanisms of cue- and context-induced seeking [55].

### Integration with other incubation mechanisms in NAcc

This study demonstrates a role for DA signaling at D1R and D2R in the NAcc in incubated cue-induced cocaine seeking, while our previous work has demonstrated that incubation depends on strengthening of glutamate transmission in the NAcc via synaptic insertion of high conductance Ca^2+^-permeable AMPA receptors (CP-AMPARs) [11,56-58]. CP-AMPAR upregulation in the NAc shell is also required for the incubation of cocaine craving [12]. This dual necessity of DA and glutamate suggests these neurotransmitters may be interacting to support incubated cue-induced seeking. There is precedent for such an interaction. For example, the reinforcing properties of the BLA-to-NAc glutamate pathway are gated by D1Rs in the NAc [59]. More generally, DA can alter how MSNs respond to glutamate signaling, although this relationship is complex and not fully understood [16,47,60]. One potentially relevant mode of interaction is suggested by the demonstration that D1R activation primes AMPAR for synaptic insertion in cultured NAc neurons [61]. Another possibility is related to the fact that MSNs *in vivo* have bi-stable resting potentials, existing in a hyperpolarized ‘down state’ and a depolarized ‘up state’ from which they can fire action potentials [62]. DA may be acting through D1R to facilitate transition to the ‘up state’ [63,64]. Alternatively, DA may indirectly increase MSN activity by reducing inhibitory tone. Activation of D2R on the D2-MSN, which collateralize on D1-MSNs, reduces GABA release from the D2-MSN resulting in a disinhibition of D1-MSN. This mechanism contributes to cocaine-induced locomotor activity [65]. Furthermore, we must consider DA actions on D2R present on cholinergic interneurons, which modulate MSN firing when inhibited via optogenetics [66].

### Methodological Considerations

There are several methodological issues that should be acknowledged. One potential concern is buffering of DA by the sensor. However, we found no difference in cocaine self-administration and incubation between rats expressing GRAB_DA2m and a mutant sensor that does not bind DA, arguing against substantial buffering. We also acknowledge that we cannot rule out a difference in DA release when comparing FAD1-2 to FAD40-50 that it is too small for our sensor to detect. In regard to pharmacology studies, the major concerns are specificity of the chosen drugs for their receptors and the possibility of off-target effects. SCH39166 was selected based on its high affinity for the D1R relative to other DA and 5-HT receptors [67,68], while L-741,626 was selected due to its 50-fold higher affinity for the D2R than the D3R [69,70]. However, we cannot completely rule out the possibility of interactions with other receptors. The principal off-target effect of DA-R antagonism is suppression of motor activity, which could reduce responding during seeking tests. To avoid this, we selected a SCH39166 dose that reduced methamphetamine seeking without eliciting motor deficits as measured by operant responding for sucrose/maltodextrin [48]. Consistent with these prior results, we found no significant reduction of locomotion during seeking tests or open field activity. For L-741,626, we likewise found no evidence of non-specific reduction of motor activity, consistent with another study that administered various doses of this drug into NAcc and reported no effects on locomotion [71]. An additional potential concern is a ‘floor effect’ in FAD1 pharmacology studies, although this concern is lessened by our use of a 1-h seeking test during which significant responding occurs even on FAD1. Our lab has successfully conducted such studies in the past [72,73].

## Conclusions

Cue-induced cocaine seeking is accompanied by a similar magnitude of NAcc DA release in early and late abstinence, suggesting that ‘incubated’ seeking does not reflect enhancement of DA release (i.e., a presynaptic effect). Instead, the suppression of incubated seeking (FAD40-50) but not FAD1 seeking) by D1R or D2R antagonists suggests that postsynaptic changes occur that enable the same amount of DA release to drive stronger behavioral responding. It will be important for future studies to examine how glutamate and DA interact postsynaptically to set the gain on cue reactivity during cocaine abstinence.

## ACKNOWLDGEMENTS

We thank Dr. John Williams for providing GRAB_DA2m virus, Dr. Rajtarun Madangopal for assistance with MedPC code and python code, Dr. Venus Sherathiya for assistance with GuPPy, and Randall Olson for assistance with MATLAB code.

## AUTHOR CONTRIBUTIONS

SJW and MEW developed the experiments and wrote the manuscript. SJW, ABK, ALM, MMB, LMK, CDM, and JGW conducted the experiments. AMW provided input on fiber photometry.

## ADDITIONAL INFORMATION

Supplementary information is provided.

