## Supplemental Material for "Dopamine transmission at D1 and D2 receptors in the nucleus accumbens contributes to the expression of incubation of cocaine craving"

**Contents**: Figures S1-S7, Tables S1-S10 (which show full statistical analyses), Supplementary Methods, and References for Supplementary Methods

### Supplementary Figures

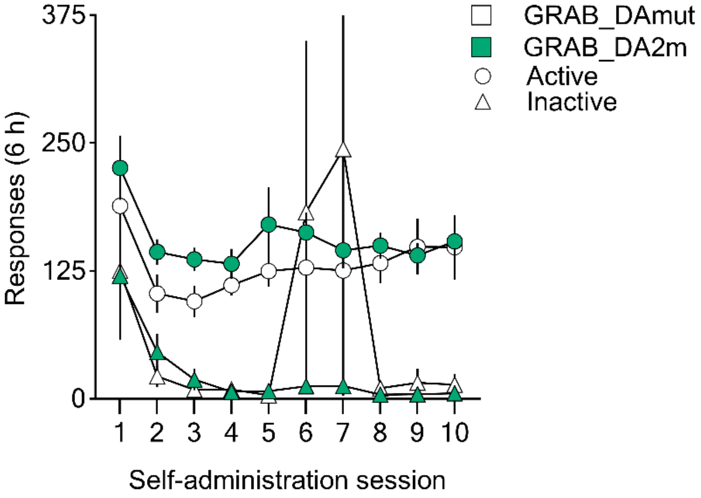


**Figure S1**. Cocaine self-administration and incubation data with no exclusions. Inactive responses for a single female rat that were excluded from Fig. 1D are included here. Rats expressing both GRAB_DA2m and GRAB_DAmut learned to nose-poke into the active port for an infusion of cocaine and discriminated between the drug-associated port and the inactive port (*p<0.05). Mean (±SEM) for each 6-h session is shown.

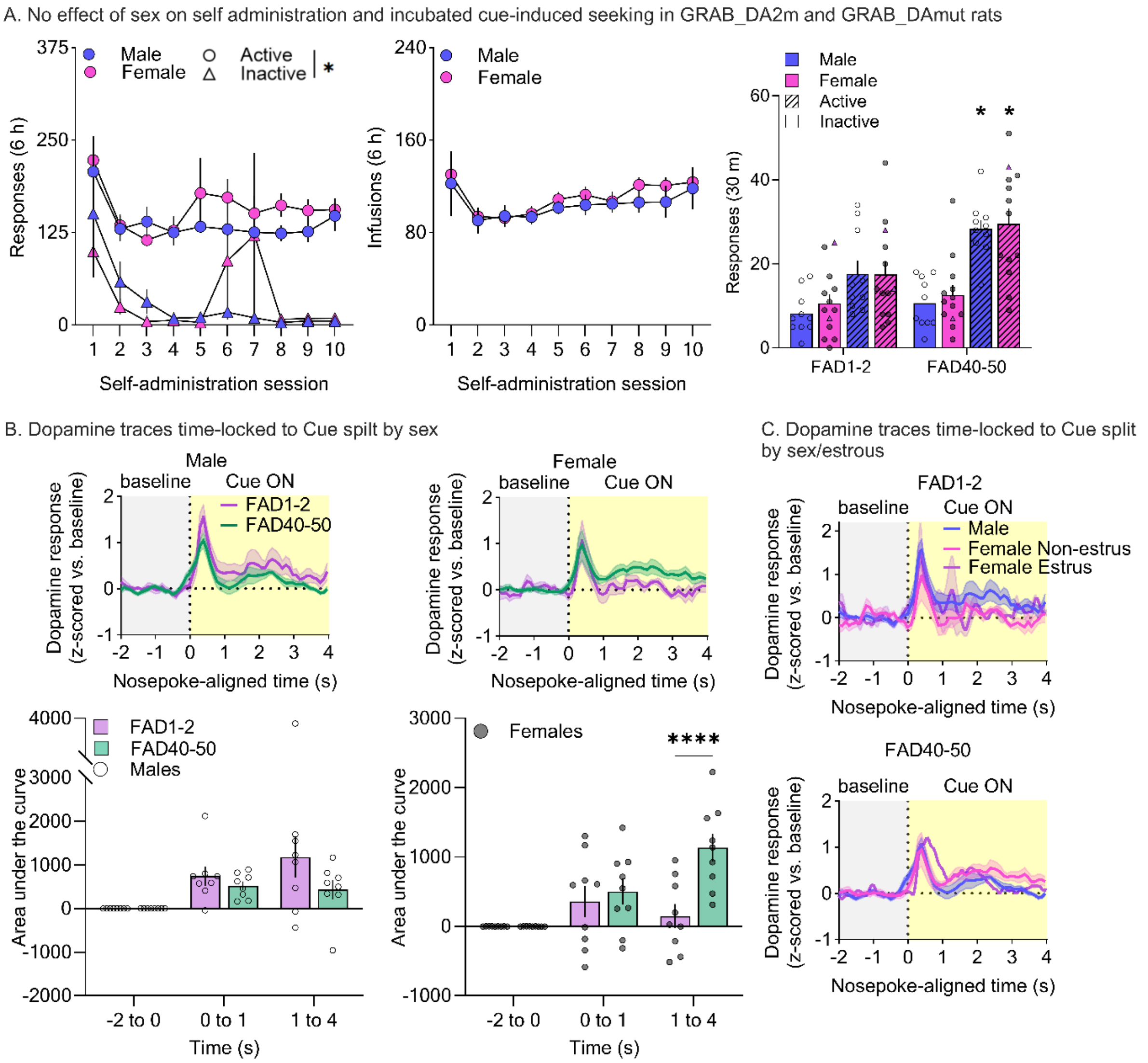


**Figure S2**. Behavioral and fiber photometry data split by sex. **A**. Self-administration and seeking test data from Fig. 1 are shown for males and females separately. *Left:* Both male and female rats expressing GRAB_DA2m or GRAB_DAmut learned to nose-poke into the active port for an infusion of cocaine and discriminated between the drug-associated port and the inactive port (*p<0.05). A mixed effect model found no main effect of sex ($F_{1,21}$=0.640, $p$=0.433; Table S3). Mean (±SEM) for each 6-h session is shown. *Middle:* Cocaine infusion data collapsed by virus group and spilt by sex. There was no main effect of sex ($F_{1,21}$=0.360, $p$=0.555; Table S3). Mean (±SEM) for each 6-h session is shown. *Right:* Each rat underwent cue-induced seeking tests on both FAD1-2 and FAD40-50 (within-subject design). Data are shown collapsed by virus and spilt by sex (GRAB_DA2m n = 8 males/9 females, GRAB_DAmut n = 2 males/4 females; *p<0.05 vs respective FAD1-2 active port data). There was no main effect of sex ($F_{1,21}$=0.296, $p$=0.5924; Table S3). Bars show mean (±SEM) for each 30-min seeking test while dots indicate individual rats. Purple triangles indicate female rats that were in estrus at the time of the seeking test (2 on FAD1-2, 1 on FAD40-50). **B**. DA traces time-locked to active pokes that triggered the cue (0-4 sec) during seeking tests depicted in **A.** Data are shown for males (*Left;* n = 8) and females (*Right;* n = 9) separately (aggregated data are shown in Fig. 2B). Traces show z-scored mean DA response normalized to a baseline period (-2 to 0 s) on FAD1-2 and FAD40-50. SEM is shown in shaded area around the mean. Bar graphs below DA traces show AUC (mean ± SEM) spilt into time epochs of baseline (-2 to 0 s), initial peak (0 to 1 s) and secondary peak (1 to 4 s). For aggregated data, we found no significant effect of sex ($F_{1,30}$=1.10, $p$=0.304) or test day ($F_{1,15}$=1.74, $p$=0.207) but a significant effect of time ($F_{2,30}$=11.5, $p$=0.000200). However, for males, there was a significant effect of time ($F_{2,14}$=6.86, $p$=0.00838) but not test day ($F_{1,7}$=2.74, $p$=0.142) and no significant interaction ($F_{2,14}$=0.451, $p$=0.0705), while females showed a significant effect of time ($F_{2,16}$=6.59, $p$=0.00827) and test day ($F_{1,8}$=12.0, $p$=0.00859) and a significant interaction ($F_{2,16}$=21.4, $p$<0.0001). Post hoc analysis of the female data found a significant increase on FAD40-50 vs FAD1-2 in DA levels 1-4 s after the cue onset ($t_{24}$=6.85, ****$p$<0.0001). Full statistical output is shown in Table S5. **C**. DA traces time-locked to active pokes that triggered the cue (0-4s) split by sex and by estrous cycle stage. *Top:* z-scored mean DA response normalized to a baseline period (-2 to 0 s) on FAD1-2 (n = 8 males, n = 7 females non-estrus, n = 2 females estrus). SEM is shown in shaded area around the mean. *Bottom:* z-scored mean DA response normalized to a baseline period (-2 to 0 s) on FAD40-50 (n = 8 males, n = 8 females non-estrus, n = 1 female estrus). FAD, forced abstinence day.

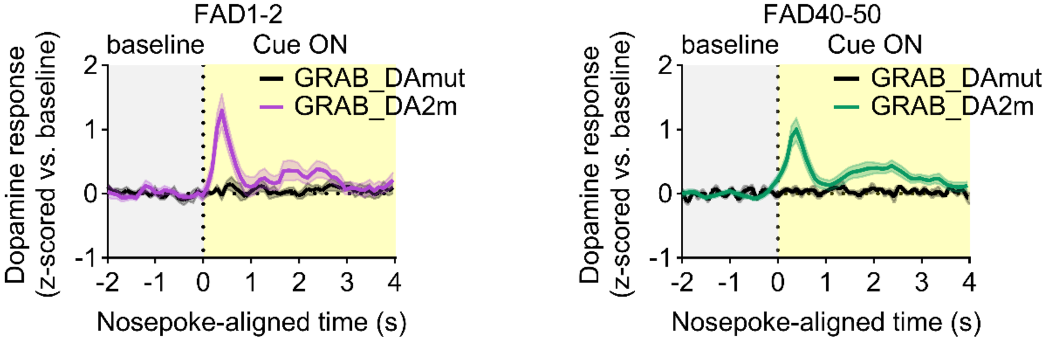


**Figure S3**. DA signals are evident in NAcc of rats expressing GRAB-DA2m but not rats expressing GRAB-DAmut. **A.** DA traces time-locked to active pokes that triggered the cue (0-4s) during the cue-induced seeking tests for GRAB_DA2m and GRAB_DAmut expressing rats. *Left:* z-scored mean DA response normalized to a baseline period (-2 to 0 s) on FAD1-2 (n = 17 GRAB_DA2m, n = 6 GRAB_DAmut). SEM is shown in shaded area around the mean. *Right:* The same analysis is shown for these rats on FAD40-50 (within-subject design).

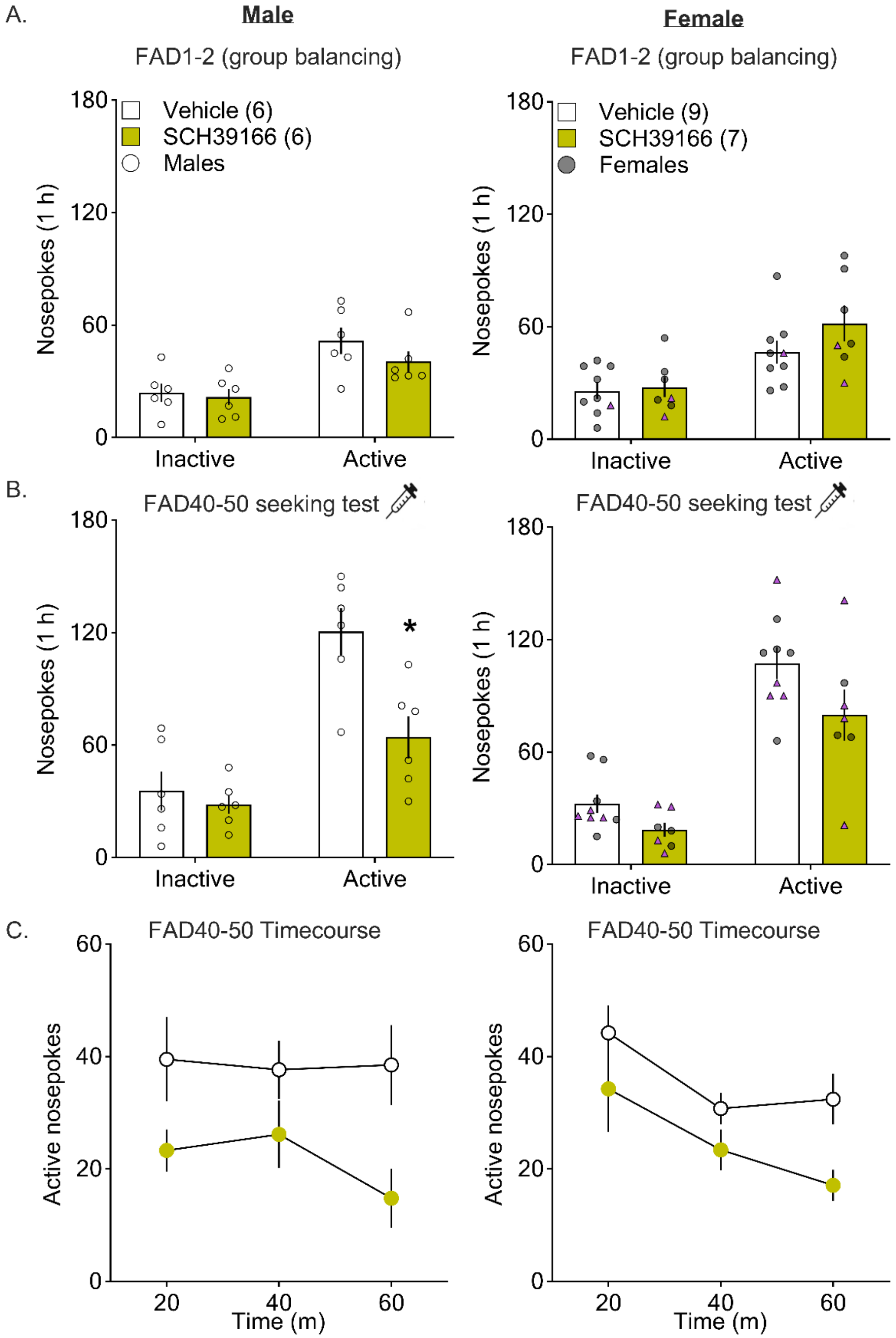
A.
Male
FAD1-2 (group balancing)
Vehicle (6)
SCH39166 (6)
Males
Nosepokes (1 h)
Inactive
Active
Female
FAD1-2 (group balancing)
Vehicle (9)
SCH39166 (7)
Females
B.
FAD40-50 seeking test
*
C.
FAD40-50 Timecourse
Active nosepokes
Time (m)

**Figure S4**. Additional analysis of SCH39166 experiment (Fig. 4) for male and female rats. Male data are shown in the left column and female data are shown in the right column. **A**. Mean (±SEM) for active port responses during FAD1-2 seeking test (no microinjection was given). These data, in combination with infusion data shown in Fig. 4D-*Left,* were used to balance groups for the FAD40-50 seeking test. **B**. Effect of intra-NAcc vehicle or SCH39166 on nose-pokes (mean ± SEM) during FAD40-50 seeking test in males and females. Aggregated data shown in Fig. 4D-*Middle* revealed significant main effects of treatment and port, and a significant treatment x port interaction, but no main effect of sex ($F_{1,22}$=0.0214, $p$=0.885) (Table S7). However, to thoroughly address sex as a biological variable, panel (B) presents further analysis of these data spilt by sex (males, n = 6 vehicle/6 SCH39166; females, n = 9 vehicle/7 SCH39166) (full statistical output in Table S8). Both males and females showed a significant main effect of treatment (vehicle vs SCH39166; male: $F_{1,10}$=8.52, $p$=0.0154; female: $F_{1,14}$=5.39, $p$=0.0358), but only males showed a significant port x treatment interaction (male: $F_{1,10}$=6.79, $p$=0.0262; female: $F_{1,14}$=0.805, $p$=0.385). Post hoc tests revealed that SCH39166 significantly reduced active port responding on FAD40-50 in males ($t_{20}$=3.91, $p$=0.00173), whereas this effect trended in females but did not achieve significance ($t_{28}$=2.34, $p$=0.0517). Combined with data in Fig. 4D, these results indicate no main effect of sex for aggregated data, although data disaggregated by sex may suggest a more robust effect in male rats of D1R blockade on incubated cocaine seeking. We did not perform this analysis on FAD1-2 data due to lower sample sizes and lack of a main effect of treatment. **C.** Mean (±SEM) pokes for FAD40-50 seeking test data shown in panel (B) split into three 20-minute bins. Dots show data for individual rats. Purple triangles indicate female rats that were in estrus at the time of the seeking test (FAD1-2: n = 2 SCH39166, n = 1 vehicle; FAD40-50: n = 4 SCH39166, n = 4 vehicle). FAD, forced abstinence day.

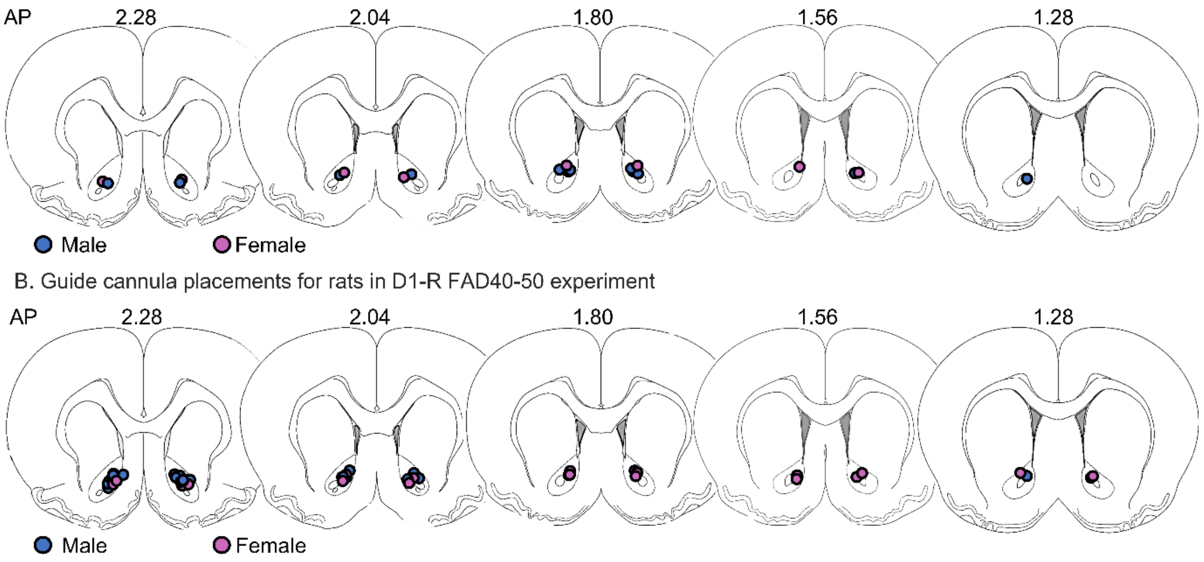


**Figure S5**. Cannula placements for experiments shown in Fig. 4. **A**. Cannula placements for FAD1 D1R antagonist experiment shown in Fig. 4B. Placement of injector tips was determined using Cresyl violet counterstaining of formalin-fixed tissue. Dots show injector tip for each cannula (bilateral) for each rat, and color indicates sex of the rat. Sections adapted from Paxinos & Watson 7$^{th}$ edition. **B**. Cannula placements for FAD40-50 D1R antagonist experiment shown in Fig. 4D. Placement of injector tips was determined as described for (A). AP, anterior posterior; FAD, forced abstinence day.

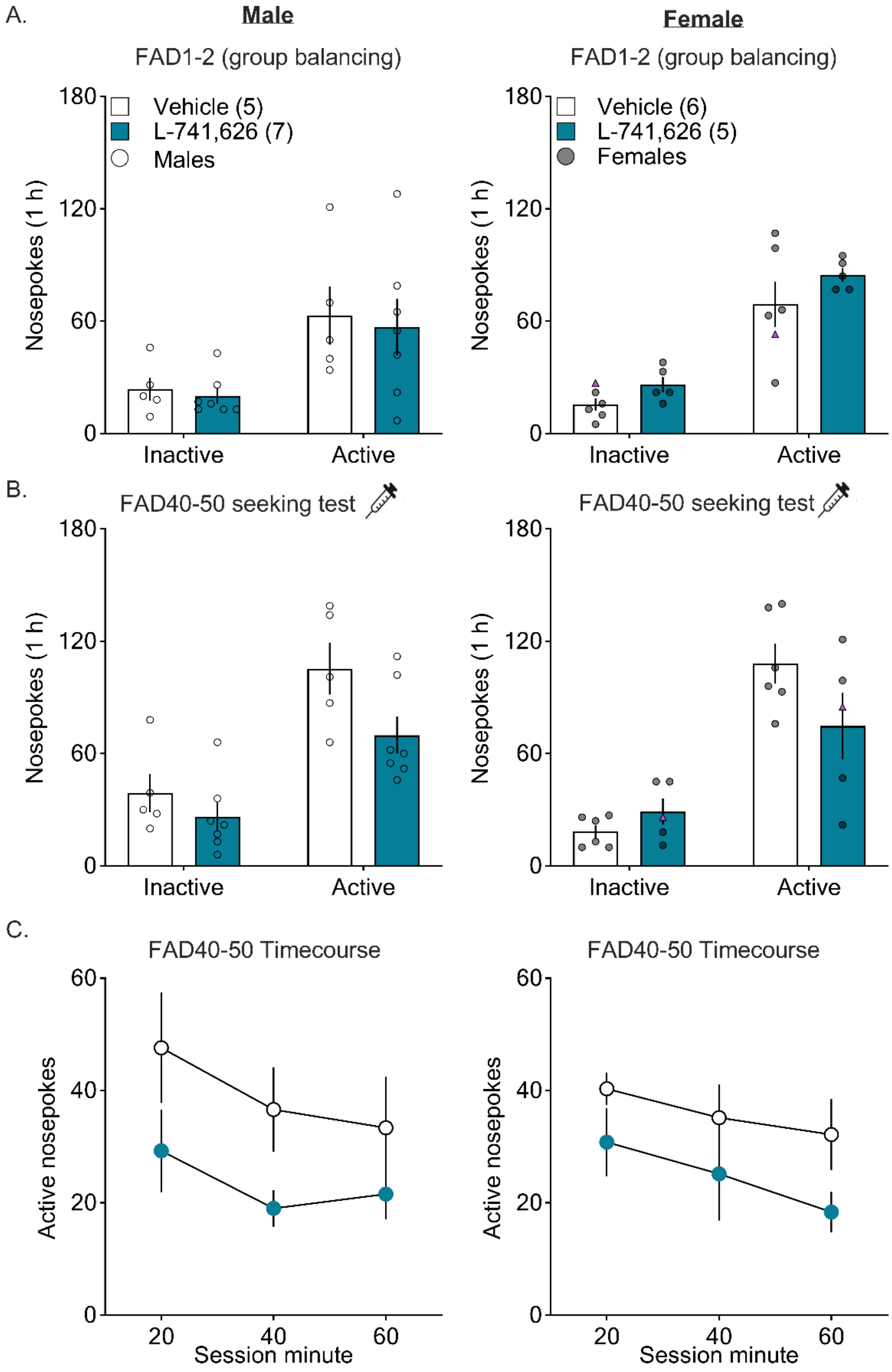


**Figure S6**. Additional analysis of L-741,626 experiment (Fig. 5) for male and female rats. Male data are shown in the left column and female data in the right column. **A.** Mean (±SEM) for active port responses during FAD1-2 seeking test (no microinjection was given). These data, in combination with infusion data shown in Fig. 5D-*Left,* were used to balance groups for the FAD40-50 seeking test. **B.** Effect of intra-NAcc vehicle or L-741,626 on nose-pokes (mean ± SEM) during FAD40-50 seeking test in males and females. Aggregated data shown in Fig. 5D-*Middle* revealed significant main effects of treatment, port, and a significant treatment x port interaction but no main effect of sex ($F_{1,19}$=0.101, $p$=0.754) (Table S9). However, to thoroughly address sex as a biological variable, panel (B) presents further analysis of these data split by sex (n = 5 males/6 females for vehicle, n = 7 males/5 females for L-741,626; full statistical output in Table S10). The male rats showed a significant effect of treatment ($F_{1,10}$=5.36, $p$=0.0432) while the females did not ($F_{1,9}$=0.839, $p$=0.384), although males failed to show a significant interaction between port and treatment ($F_{1,10}$=1.26, $p$=0.289). Combined with data in Fig. 5D, these results indicate no main effect of sex for aggregated data, although data disaggregated by sex may suggest a more robust effect in male rats of D2R blockade on incubated seeking. We did not perform this analysis on FAD1-2 data due to lower sample sizes and lack of a main effect of treatment. **C.** Mean (±SEM) pokes for FAD40-50 seeking test data spilt into three 20-min bins. Dots show data for individual rats. Purple triangles indicate female rats that were in estrus at the time of the seeking test (FAD1-2: n = 0 L-741,626, n = 1 vehicle; FAD40-50: n = 1 L-741,626, n = 0 vehicle). FAD, forced abstinence day.

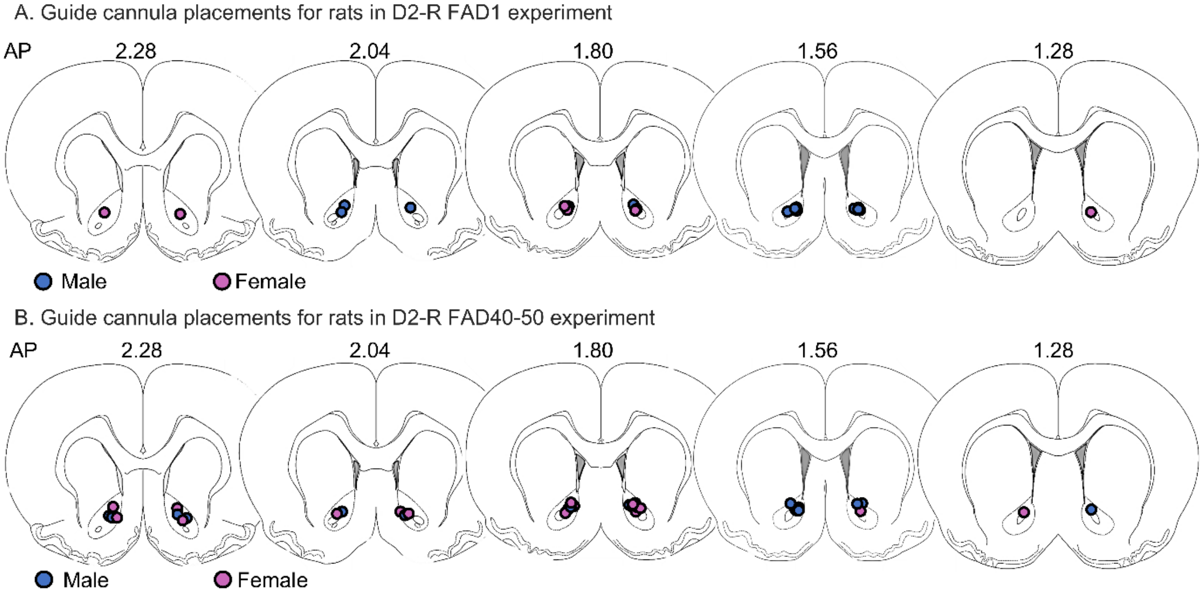


**Figure S7**. Cannula placements for experiments shown in Fig. 5. **A**. Cannula placements for FAD1 D2R antagonist experiments. Placement of injector tips was determined using Cresyl violet counterstaining of formalin-fixed tissue. Dots show injector tip for each cannula (bilateral) for each rat, and color indicates sex of the rat. Sections adapted from Paxinos & Watson 7th edition. **B**. Cannula placements for FAD40-50 D2R antagonist experiments. Placement of injector tips was determined as described for (A). AP, anterior posterior; FAD, forced abstinence day.

### Supplementary Tables

#### Table S1. Number of subjects that successfully reached different phases of each experiment

| **Figs** | **Group** | **Self-Administration** | **Pre-screening** | **Seeking test** | **Animals included after histology** |
|---|---|---|---|---|---|
| **1-3** | GRAB_DAmut | 7 (3M, 4F) | 7 (3M, 4F) | 7 (3M, 4F) | 6 (2M, 4F) |
| | GRAB_DA2m | 28 (13M, 15F) | 25 (12M, 13F) | 23 (11M,12F) | 17 (8M, 9F) |
| **Figs** | **Group** | **Self-Administration** | **Seeking test** | | **Animals included after histology** |
| **4** | FAD1-2 | 16 (8M, 8F) | 14 (8M, 6F) | | 13 (7M, 6F) |
| | FAD40-50 | 40 (18M,22F) | 39 (17M,22F) | | 28 (12M, 16F) |
| **5** | FAD1-2 | 16 (7M, 9F) | 15 (6M, 9F) | | 13 (5M, 8F) |
| | FAD40-50 | 34 (19M, 15F) | 38 (18M, 15F) | | 23 (12M, 11F) * |
| **Figs** | **Group** | **Open Field testing** | | | **Animals included after histology** |
| **4-5** | Open field D1 | 8 M | | | 7 M |
| | Open field D2 | 6 (3M, 3F) | | | 5 (3M, 2F) |
| **Total** | | 155 (79M, 76F) from surgery | | | 112 (56M, 56F) |

*7 rats were excluded due to pump failure during microinjection

F, female; M, male

**Table S2. Statistical output for immunohistochemical and behavioral data in Figs. 1 and 3**

| Expt phase | Measure | Fixed effects in model | F-value | P-value | Significant? | Figure |
|---|---|---|---|---|---|---|
| N/A | Fluorescence Intensity | Group | $F_{2,11}$=10.97 | 0.0024 | ** | 1B |
| | | Holm-Šídák†† | | | | |
| | | No virus vs. 3 weeks | $t_{11}$=4.419 | 0.0031 | ** | |
| | | No virus vs. 10 weeks | $t_{11}$=4.341 | 0.0031 | ** | |
| | | 3 weeks vs. 10 weeks | $t_{11}$=0.1095 | 0.915 | n.s. | |
| SA | Nose pokes over 6-h (n=17 GRAB_DA2m, 6 GRAB_DAmut) | Virus x Session x Port | | | | 1D, left |
| | | Virus | $F_{1,189}$=0.1536 | 0.6955 | n.s. | |
| | | Session | $F_{9,189}$=3.93 | 0.0001 | *** | |
| | | Port | $F_{1,21}$=37.4 | <0.0001 | **** | |
| | | Virus x Session | $F_{9,189}$=1.726 | 0.0856 | n.s. | |
| | | Virus x Port | $F_{1,189}$=3.991 | 0.0472 | * | |
| | | Session x Port | $F_{9,189}$=2.106 | 0.0309 | * | |
| | | Virus x Session x Port | $F_{9,189}$=2.040 | 0.0371 | * | |
| SA | Infusions over 6-h (n=17 GRAB_DA2m, 6 GRAB_DAmut) | Virus x Session | | | | 1D, middle |
| | | Virus | $F_{1,21}$=0.2480 | 0.6237 | n.s. | |
| | | Session | $F_{9,189}$=4.172 | <0.0001 | **** | |
| | | Virus x Session | $F_{9,189}$=1.101 | 0.3641 | n.s. | |
| Cocaine Seeking | Nose pokes over 30-m (n=17 GRAB_DA2m, 6 GRAB_DAmut) | Virus x Test Day x Port | | | | 1D, right |
| | | Virus | $F_{1,21}$=0.3913 | 0.5384 | n.s. | |
| | | Test Day | $F_{1,21}$=7.526 | 0.0122 | * | |
| | | Port | $F_{1,21}$=50.05 | <0.0001 | **** | |
| | | Virus x Test Day | $F_{1,21}$=0.1149 | 0.7380 | n.s. | |
| | | Virus x Port | $F_{1,21}$=0.04717 | 0.8320 | n.s. | |
| | | Test Day x Port | $F_{1,21}$=9.071 | 0.00664 | ** | |
| | | Virus x Session x Port | $F_{1,21}$=0.04448 | 0.8350 | n.s. | |
| Cocaine Seeking | Nose pokes over 30-m (n=10M, 13F) | Test Day x Bin† | | | | 3B |
| | | Test Day | $F_{1,17}$=10.7 | 0.0001 | *** | |
| | | Bin | $F_{2,34}$=12.02 | 0.0045 | ** | |
| | | Test Day x Bin | $F_{2,34}$=0.6467 | 0.5301 | n.s. | |

†Random effects with SD = 0 excluded from model
††adjusted P-values reported for post hoc comparisons
*p<0.05, **p<0.01, ***p<0.001, ****p<0.0001
F, female; M, male

**Table S3. Statistical output for behavioral data in Fig. S2**

| Expt phase | Measure | Fixed effects in model | F-value | P-value | Significant? | Figure |
|---|---|---|---|---|---|---|
| SA | Nose pokes over 6-h (n=10M, 13F) | Sex x Session x Port | | | | S2A, left |
| | | Sex | $F_{1,21}$=0.6399 | 0.4327 | n.s. | |
| | | Session | $F_{9,189}$=3.845 | 0.0002 | *** | |
| | | Port | $F_{1,21}$=53.85 | <0.0001 | **** | |
| | | Sex x Session | $F_{9,189}$=1.002 | 0.4397 | n.s. | |
| | | Sex x Port | $F_{1,21}$=0.1336 | 0.7184 | n.s. | |
| | | Session x Port | $F_{9,189}$=0.8727 | 0.5507 | n.s. | |
| | | Sex x Session x Port | $F_{9,189}$=0.6181 | 0.7808 | n.s. | |
| SA | Infusions over 6-h (n=10M, 13F) | Sex x Session | | | | S2A, middle |
| | | Sex | $F_{1,21}$=0.3597 | 0.5551 | n.s. | |
| | | Session | $F_{9,189}$=4.188 | <0.0001 | ***** | |
| | | Sex x Session | $F_{9,189}$=0.2139 | 0.9922 | n.s. | |
| Cocaine Seeking | Nose pokes over 30-m (n=10M, 13F) | Sex x Test Day x Port | | | | S2A, right |
| | | Sex | $F_{1,21}$=0.2955 | 0.5924 | n.s. | |
| | | Test Day | $F_{1,21}$=10.62 | 0.0038 | ** | |
| | | Port | $F_{1,21}$=63.60 | 0.0001 | **** | |
| | | Sex x Test Day | $F_{1,21}$=0.01097 | 0.9176 | n.s. | |
| | | Sex x Port | $F_{1,21}$=0.2576 | 0.6171 | n.s. | |
| | | Test Day x Port | $F_{1,21}$=12.07 | 0.0023 | ** | |
| | | Sex x Test Day x Port | $F_{1,21}$=0.1273 | 0.7248 | n.s. | |

F, female; M, male
**p<0.01, ***p<0.001, ****p<0.0001

**Table S4. Statistical output for fiber photometry data in Fig. 2**

| Expt phase | Measure | Fixed effects in model | F-value | P-value | Significant? | Figure |
|---|---|---|---|---|---|---|
| Cocaine seeking (30-min) | DA response to active nose pokes that trigger cue, AUC (n=17) | Time x Test Day | | | | 2C, left |
| | | Time | $F_{2,32}$=14.0 | <0.0001 | **** | |
| | | Test Day | $F_{1,16}$=0.1281 | 0.7251 | n.s. | |
| | | Time x Test Day | $F_{2,32}$=0.4512 | 0.6409 | n.s. | |
| Cocaine seeking (30-min) | DA response to active nose pokes during cue, AUC (n=15 FAD1-2, n=17 FAD40-50) | Time x Test Day† | | | | 2C, middle |
| | | Time | $F_{2,32}$=1.361 | 0.2709 | n.s. | |
| | | Test Day | $F_{1,16}$=3.312 | 0.0875 | n.s. | |
| | | Time x Test Day | $F_{2,32}$=1.266 | 0.2987 | n.s. | |
| Cocaine seeking (30-min) | DA response to inactive nose pokes, AUC (n=17) | Time x Test Day | | | | 2C, right |
| | | Time | $F_{2,32}$=6.97 | 0.00307 | ** | |
| | | Test Day | $F_{1,16}$=1.020 | 0.3275 | n.s. | |
| | | Time x Test Day | $F_{2,32}$=2.802 | 0.0756 | n.s. | |
| Cocaine seeking (30-min) | Frequency of events | Paired t-test | | | | 2E, left |
| | | Test Day | $t_{16}$=0.7640 | 0.4388 | n.s. | |
| Cocaine seeking (30-min) | Amplitude of events | Paired t-test | | | | 2E, right |
| | | Test Day | $t_{16}$=2.84 | 0.0117 | * | |

†Random effects with SD = 0 excluded from model
*p<0.05, **p<0.01, ****p<0.0001
F, female; M, male

**Table S5. Statistical output for fiber photometry data related to Figs. S2 and 3E**

| Expt phase | Measure | Fixed effects in model | F-value | P-value | Significant? | Figure |
|---|---|---|---|---|---|---|
| Cocaine seeking (30-min) | DA response to active nose pokes that trigger cue, AUC (n=17) | Time x Sex x Test Day | | | | Data from 2C left, re-analyzed with sex as a factor |
| | | Time | $F_{2,30}$=11.46 | 0.0002 | *** | |
| | | Sex | $F_{1,30}$=1.10 | 0.304 | n.s. | |
| | | Test Day | $F_{1,15}$=1.740 | 0.2069 | n.s. | |
| | | Time x Sex | $F_{2,30}$=0.6153 | 0.5472 | n.s. | |
| | | Time x Test Day | $F_{2,30}$=1.417 | 0.2582 | n.s. | |
| | | Sex x Test Day | $F_{1,30}$=2.110 | 0.1567 | n.s. | |
| | | Time x Sex x Test Day | $F_{2,30}$=2.789 | 0.0775 | n.s. | |
| Cocaine seeking (30-min) | DA response to active nose pokes that trigger cue, AUC (n=8M) | Time x Test Day | | | | S2B, bottom left |
| | | Time | $F_{2,14}$=6.862 | 0.0084 | ** | |
| | | Test Day | $F_{1,7}$=2.738 | 0.1420 | n.s. | |
| | | Time x Test Day | $F_{2,14}$=0.4512 | 0.0705 | n.s. | |
| Cocaine seeking (30-min) | DA response to active nose pokes that trigger cue, AUC (n=9F) | Time x Test Day | | | | S2B, bottom right |
| | | Time | $F_{2,16}$=6.588 | 0.0083 | ** | |
| | | Test Day | $F_{1,8}$=11.96 | 0.0086 | ** | |
| | | Time x Test Day | $F_{2,16}$=21.37 | <0.0001 | **** | |
| | | Holm-Šídák: FAD1-2 vs. FAD40-50†† | | | | |
| | | -2 to 0 | $t_{24}$=0.001099 | 0.9991 | n.s. | |
| | | 0 to 1 | $t_{24}$=1.011 | 0.3223 | n.s. | |
| | | 1 to 4 | $t_{24}$=6.848 | <0.0001 | **** | |
| | | Holm-Šídák: FAD1-2†† | | | | |
| | | -2 to 0 vs. 0 to 1 | $t_{32}$=1.802 | 0.2238 | * | |
| | | -2 to 0 vs. 1 to 4 | $t_{32}$=0.7506 | 0.5114 | *** | |
| | | 0 to 1 vs. 1 to 4 | $t_{32}$=1.051 | 0.5114 | n.s. | |
| | | Holm-Šídák: FAD40-50†† | | | | |
| | | -2 to 0 vs. 0 to 1 | $t_{32}$=2.538 | 0.0162 | * | |
| | | -2 to 0 vs. 1 to 4 | $t_{32}$=5.733 | <0.0001 | **** | |
| | | 0 to 1 vs. 1 to 4 | $t_{32}$=3.195 | 0.0063 | ** | |
| Cocaine seeking | DA response to 1st active nose poke that triggered cue, AUC (n=17) | Time x Test Day† | | | | AUC data from traces in Fig. 3E |
| | | Time | $F_{2,32}$=3.31 | 0.0496 | * | |
| | | Test Day | $F_{1,16}$=0.5601 | 0.4651 | n.s. | |
| | | Time x Test Day | $F_{2,32}$=0.2274 | 0.7979 | n.s. | |

F, female; M, male

*p<0.05, **p<0.01, ***p<0.001, ****p<0.0001

### Table S6. Statistical output for bootstrapping analyses in Fig. 2

| Expt phase | Measure | Factors in analysis | Time 95% CI ≠ 0 | FAD1-2 vs FAD40-50 | Figure |
|---|---|---|---|---|---|
| Cocaine seeking (30-min) | DA response to active nose pokes that trigger cue, z-scored trace (n=17) | Bootstrapping<br>FAD1-2 | 0.14 – 0.76 s<br>1.61 – 2.85 s | n.s. | 2B, left |
| | | FAD40-50 | -0.59 – -0.46 s<br>-0.12 – 0.92 s<br>1.2 – 3.6 s | | |
| Cocaine seeking (30-min) | DA response to active nose pokes during cue, z-scored trace (n=15 FAD1-2, n=17 FAD40-50) | Bootstrapping<br>FAD1-2 | -1.26 – -1.13 s<br>0.29 – 0.54 s | n.s. | 2B, middle |
| | | FAD40-50 | 0.05 – 0.44 s<br>2.62 – 3.44 s<br>3.47 – 4.0 s | | |
| Cocaine seeking (30-min) | DA response to inactive nose pokes, z-scored trace (n=17) | Bootstrapping<br>FAD1-2 | 1.62 – 1.80 s<br>1.93 – 2.22 s | Yes, 1.93 – 2.22 s | 2B, right |
| | | FAD40-50 | 0.17 – 0.36 s<br>2.97 – 3.23 s | | |

**Table S7. Statistical output for D1R antagonist experiments in Fig. 4**

| Expt phase | Measure | Fixed effects in model | F-value | P-value | Significant? | Figure |
|---|---|---|---|---|---|---|
| Cocaine Seeking FAD1 | Nose pokes over 1-h (n=6 SCH39166, n=7 vehicle) | Treatment x Port | | | | 4B, middle |
| | | Treatment | $F_{1,11}$=0.1495 | 0.7064 | n.s. | |
| | | Port | $F_{1,11}$=24.54 | 0.0004 | *** | |
| | | Port x Treatment | $F_{1,11}$=0.001385 | 0.9710 | n.s. | |
| Cocaine Seeking FAD1 | Nose pokes in 20-m bins (n=6 SCH39166, 7 vehicle) | Treatment x Bin | | | | 4B, right |
| | | Treatment | $F_{1,12}$=0.1543 | 0.7014 | n.s. | |
| | | Bin | $F_{2,24}$=5.21 | 0.0132 | * | |
| | | Bin x Treatment | $F_{2,24}$=0.03536 | 0.9653 | n.s. | |
| Cocaine Seeking FAD40-50 | Nose pokes over 1-h (n=13 SCH39166, 15 vehicle) | Treatment x Port | | | | 4D, middle |
| | | Treatment | $F_{1,26}$=14.13 | 0.0009 | *** | |
| | | Port | $F_{1,26}$=115.6 | <0.0001 | **** | |
| | | Port x Treatment | $F_{1,26}$=6.03 | 0.0210 | * | |
| | | Holm-Šídák†† | | | | |
| | | Inactive | $t_{52}$=1.182 | 0.2425 | n.s. | |
| | | Active | $t_{52}$=4.44 | <0.0001 | *** | |
| Cocaine Seeking FAD40-50 | Nose pokes in 20-m bins (n=13 SCH39166, 15 vehicle) | Treatment x Bin | | | | 4D, right |
| | | Treatment | $F_{1,26}$=16.7 | 0.0124 | ** | |
| | | Bin | $F_{2,52}$=4.79 | 0.000377 | ** | |
| | | Bin x Treatment | $F_{2,52}$=1.090 | 0.3438 | n.s. | |
| Cocaine Seeking FAD1 | Beambreaks over 1-h (n=6 SCH39166, 7 vehicle) | Unpaired t test | | | | 4E, left |
| | | Treatment | $t_{10}$=2.43 | 0.0352 | * | |
| Cocaine Seeking FAD40-50 | Beambreaks over 1-h (n=13 SCH39166, 15 vehicle) | Unpaired t test | | | | 4E, middle |
| | | Treatment | $t_{24}$=0.6012 | 0.5533 | n.s. | |
| Open Field | Distance traveled (n=7M) | Unpaired t test | | | | 4E, right |
| | | Treatment | $t_7$=0.3153 | 0.7617 | n.s. | |

††adjusted P-values reported for post hoc comparisons
*p<0.05, **p<0.01, ***p<0.001, ****p<0.0001
Analysis shown for combined female (F) and male (M) data.

**Table S8. Statistical output for D1R antagonist experiments related to Fig. S4**

| Expt phase | Measure | Fixed effects in model | F-value | P-value | Significant? | Figure |
|---|---|---|---|---|---|---|
| Cocaine Seeking FAD40-50 | Nose pokes over 1-h (n=13 SCH39166, 15 vehicle) | Sex x Treatment x Port | | | | Data from 4D middle, re-analyzed with sex as a factor |
| | | Sex | $F_{1,22}$=0.02136 | 0.8851 | n.s. | |
| | | Treatment | $F_{1,22}$=14.46 | 0.0010 | *** | |
| | | Port | $F_{1,22}$=104.8 | <0.0001 | **** | |
| | | Sex x Treatment | $F_{1,22}$=0.2906 | 0.5952 | n.s. | |
| | | Sex x Port | $F_{1,22}$=0.5476 | 0.4671 | n.s. | |
| | | Treatment x Port | $F_{1,22}$=6.766 | 0.0163 | * | |
| | | Sex x Treatment x Port | $F_{1,22}$=1.549 | 0.2264 | n.s. | |
| Cocaine Seeking FAD40-50 | Nose pokes over 1-h (n=6M SCH39166, 6M vehicle) | Treatment x Port | | | | S4B, left |
| | | Treatment | $F_{1,10}$=8.515 | 0.0154 | * | |
| | | Port | $F_{1,10}$=41.41 | <0.0001 | **** | |
| | | Port x Treatment | $F_{1,10}$=6.791 | 0.0262 | * | |
| | | Holm-Šídák†† | | | | |
| | | Inactive | $t_{20}$=0.5092 | 0.6162 | n.s. | |
| | | Active | $t_{20}$=3.912 | 0.0017 | ** | |
| Cocaine Seeking FAD40-50 | Nose pokes in 20-m bins (n=6M SCH39166, 6M vehicle) | Treatment x Bin | | | | S4C, left |
| | | Treatment | $F_{1,10}$=10.16 | 0.0097 | ** | |
| | | Bin | $F_{2,20}$=0.5330 | 0.5949 | n.s. | |
| | | Bin x Treatment | $F_{2,20}$=0.5980 | 0.5595 | n.s. | |
| Cocaine Seeking FAD40-50 | Nose pokes over 1-h (n=7F SCH39166, 9F vehicle) | Treatment x Port | | | | S4A, right |
| | | Treatment | $F_{1,14}$=5.391 | 0.0358 | * | |
| | | Port | $F_{1,14}$=79.50 | <0.0001 | **** | |
| | | Port x Treatment | $F_{1,14}$=0.8050 | 0.3848 | n.s. | |
| Cocaine Seeking FAD40-50 | Nose pokes in 20-m bins (n=7F SCH39166, 9F vehicle) | Treatment x Bin | | | | S4C, right |
| | | Treatment | $F_{1,14}$=6.067 | 0.0273 | * | |
| | | Bin | $F_{2,28}$=6.850 | 0.0038 | ** | |
| | | Bin x Treatment | $F_{2,28}$=0.4669 | 0.6317 | n.s. | |

††adjusted P-values reported for post hoc comparisons
*p<0.05, **p<0.01, ***p<0.001, ****p<0.0001
F, female; M, male

### Table S9. Statistical output for D2R antagonist experiments in Fig. 5

| Expt phase | Measure | Fixed effects in model | F-value | P-value | Significant? | Figure |
|---|---|---|---|---|---|---|
| Cocaine Seeking FAD1 | Nose pokes over 1-h (n=6 L-741,626, 7 vehicle) | Treatment x Port[†] | | | | 5B, middle |
| | | Treatment | $F_{1,22}$=0.4189 | 0.5242 | n.s. | |
| | | Port | $F_{1,22}$=12.17 | 0.0021 | ** | |
| | | Port x Treatment | $F_{1,22}$=0.8257 | 0.3734 | n.s. | |
| Cocaine Seeking FAD1 | Nose pokes in 20-m bins (n=6 L-741,626, 7 vehicle) | Treatment x Bin | | | | 5B, right |
| | | Treatment | $F_{1,13}$=0.07718 | 0.7855 | n.s. | |
| | | Bin | $F_{2,26}$=13.31 | 0.0001 | *** | |
| | | Bin x Treatment | $F_{2,26}$=0.2231 | 0.8015 | n.s. | |
| Cocaine Seeking FAD40-50 | Nose pokes over 1-h (n=12 L-741,626, 11 vehicle) | Treatment x Port | | | | 5D, middle |
| | | Treatment | $F_{1,21}$=5.217 | 0.0329 | * | |
| | | Port | $F_{1,21}$=89.72 | <0.0001 | **** | |
| | | Port x Treatment | $F_{1,21}$=7.06 | 0.0148 | * | |
| | | Holm-Šídák[††] | | | | |
| | | Inactive | $t_{42}$=0.03070 | 0.9757 | n.s. | |
| | | Active | $t_{42}$=3.46 | 0.00251 | ** | |
| Cocaine Seeking FAD40-50 | Nose pokes in 20-m bins (n=12 L-741,626, 11 vehicle) | Treatment x Bin | | | | 5D, right |
| | | Treatment | $F_{1,21}$=6.32 | 0.0202 | * | |
| | | Bin | $F_{2,42}$=5.41 | 0.00811 | ** | |
| | | Bin x Treatment | $F_{2,42}$=0.4008 | 0.6723 | n.s. | |
| Cocaine Seeking FAD1 | Beambreaks over 1-h (n=6 L-741,626, 7 vehicle) | Unpaired t test | | | | 5E, left |
| | | Treatment | $t_{11}$=0.7417 | 0.4738 | n.s. | |
| Cocaine Seeking FAD40-50 | Beambreaks over 1-h (n=12 L-741,626, 11 vehicle) | Unpaired t test | | | | 5E, middle |
| | | Treatment | $t_{20}$=1.950 | 0.0654 | n.s. | |
| Open Field | Distance traveled (n= 2F, 3M) | Unpaired t test | | | | 5E, right |
| | | Treatment | $t_4$=0.5496 | 0.6118 | n.s. | |

Analysis shown for combined female (F) and male (M) data.
*p<0.05, **p<0.01, ***p<0.001, ****p<0.0001

**Table S10. Statistical output for D2R antagonist experiments related to Fig. S6**

| Expt phase | Measure | Fixed effects in model | F-value | P-value | Significant? | Figure |
|---|---|---|---|---|---|---|
| Cocaine Seeking FAD40-50 | Nose pokes over 1-h (n=12 L-741,626, 11 vehicle) | <u>Sex x Treatment x Port</u> | | | | Data from 5D middle, re-analyzed with sex as a factor |
| | | Sex | $F_{1,19}$=0.1012 | 0.7538 | n.s. | |
| | | Treatment | $F_{1,19}$=4.859 | 0.0400 | * | |
| | | Port | $F_{1,19}$=84.85 | <0.0001 | **** | |
| | | Sex x Treatment | $F_{1,19}$=0.6303 | 0.4370 | n.s. | |
| | | Sex x Port | $F_{1,19}$=0.9264 | 0.3479 | n.s. | |
| | | Treatment x Port | $F_{1,19}$=6.289 | 0.0214 | * | |
| | | Sex x Treatment x Port | $F_{1,19}$=0.6325 | 0.4363 | n.s. | |
| Cocaine Seeking FAD40-50 | Nose pokes over 1-h (n=7M L-741,626, 5M vehicle) | <u>Treatment x Port</u> | | | | S6B, left |
| | | Treatment | $F_{1,10}$=5.358 | 0.0432 | * | |
| | | Port | $F_{1,10}$=29.15 | 0.0003 | *** | |
| | | Port x Treatment | $F_{1,10}$=1.256 | 0.2885 | n.s. | |
| Cocaine Seeking FAD40-50 | Nose pokes over 20-m (n=7M L-741,626, 5M vehicle) | <u>Treatment x Bin</u> | | | | S6C, left |
| | | Treatment | $F_{1,10}$=4.048 | 0.0719 | n.s. | |
| | | Bin | $F_{2,20}$=3.154 | 0.0645 | n.s. | |
| | | Bin x Treatment | $F_{2,20}$=0.2563 | 0.7764 | n.s. | |
| Cocaine Seeking FAD40-50 | Nose pokes over 1-h (n= 5F L-741,626, 6F vehicle) | <u>Treatment x Port</u> | | | | S6B, right |
| | | Treatment | $F_{1,9}$=0.8392 | 0.3835 | n.s. | |
| | | Port | $F_{1,9}$=65.09 | <0.0001 | **** | |
| | | Port x Treatment | $F_{1,9}$=6.860 | 0.0278 | * | |
| | | <u>Holm-Šídák</u>†† | | | | |
| | | Inactive | $t_{18}$=0.7124 | 0.4853 | n.s. | |
| | | Active | $t_{18}$=2.229 | 0.0762 | n.s. | |
| Cocaine Seeking FAD40-50 | Nose pokes over 20-m (n= 5F L-741,626, 6F vehicle) | <u>Treatment x Bin</u> | | | | S6C, right |
| | | Treatment | $F_{1,9}$=2.804 | 0.1284 | n.s. | |
| | | Bin | $F_{2,18}$=3095 | 0.0699 | n.s. | |
| | | Bin x Treatment | $F_{2,18}$=0.1587 | 0.8545 | n.s. | |

†Random effects with SD = 0 excluded from model
††adjusted P-values reported for post hoc comparisons
*p<0.05, ***p<0.001, ****p<0.0001
F, female; M, male

### Supplementary Methods

#### Subjects

We used male and female Long-Evans rats, either obtained from Charles River (Wilmington, MA) or bred in house. Rats received from Charles River were ~9 weeks upon receipt and were group housed for ~1 week after arrival to acclimate prior to surgery. Rats bred in house were group housed with same sex litter mates from weaning until surgery at 10-17 weeks of age. After surgery, all rats were single housed. Throughout the experiment, rats had free access to standard laboratory chow and water in their home cages, and were maintained on a reverse 12:12h light/dark cycle (lights off at 10 AM prior to surgery and 9-9:30 AM post-surgery due to a switch in housing rooms after surgery). All procedures were approved by the OHSU Animal Care and Use Committee and followed NIH guidelines outlined in the Guide for the Care and Use of Laboratory Animals. From a total of 79 male and 76 female rats assigned to experimental groups, 10 were excluded due to failure to acquire self-administration, equipment failure during testing (n = 7), significant outlier values during behavioral testing (n = 2), or unexpected deaths during forced abstinence (n = 2). After histology, 32 were excluded due to misplaced intracranial cannula and 1 due to signs of infection surrounding the cannula shaft. See Table S1 for specifics.

#### Drugs

We received cocaine HCl from the NIDA Drug Supply Program. Cocaine was dissolved in sterile 0.9% saline and buffered with NaOH to reach pH 7. The final concentration of the stock solution was 5 mg/mL. Cocaine was stored at 4˚C once in solution. We dissolved the dopamine (DA) D1 receptor selective antagonist SCH39166 hydrobromide (Tocris Bioscience, Cat. #2299) in DMSO (Sigma Aldrich, #276855) at 500 mM. We then diluted 1:10 in sterile saline to make 50 mM stock vials that were stored at -20ºC. On the day of the experiment, this stock was diluted 1:10 in sterile saline to make a 5 mM working stock solution; 0.5 µl was injected into each hemisphere, delivering 1 µg SCH 39166/hemisphere. This dose was based on efficacy in prior intra-NAcc infusion studies [1]. As the SCH39166 working stock was 1% DMSO in sterile saline, we used 1% DMSO in sterile saline in the vehicle group for SCH39166 experiments. We dissolved the DA D2 receptor selective antagonist L-741,626 (Tocris Bioscience, Cat. #1003/10) in 200 proof ethanol (Decon Laboratories, Inc; #2701) at 100 mM, diluted 1:2 with Tween-80 (Sigma, P4780), and finally diluted 1:10 with sterile water for a final working solution of 5 mM L-741,626. Working solutions were stored at -20ºC and slow thawed/vortexed the day of testing. Accordingly, we used a solution of 5% ethanol, 5% Tween-80, and 90% sterile water for vehicle groups in L-741,626 experiments. This vehicle and dose were based on other intra-NAcc studies [2].

#### Surgery

Surgical procedures depended on whether rats were destined for: 1) cocaine self-administration (SA) followed by fiber photometry measurements of DA levels in NAcc during cue-induced seeking tests, 2) cocaine SA followed by intracranial infusion of DA antagonists prior to cue-induced seeking tests, or 3) intracranial DA receptor antagonist infusion into nucleus accumbens core (NAcc) prior to open field experiments.

*Intravenous catheter implantation:* Rats destined for cocaine SA experiments were implanted with silastic catheters as previously described [3,4]. Briefly, rats were anesthetized with inhalable isoflurane (5% for induction, 1-3% maintenance) and a catheter (Plastics One part #313000BM-15 with Liveo Laboratory Tubing #508-001) was passed subcutaneously from the mid-scapular region and inserted into the right jugular vein. The external portion of the catheter is attached to a mesh back mount platform to which an infusion line can be connected during intravenous drug SA.

*Intracranial cannula implantation:* For rats receiving intracranial drug infusions after drug SA, intravenous catheterization was immediately followed by intracranial surgery to implant bilateral 23-gauge guide cannula (Plastics One C317G) above the NAcc. For intra-NAcc drug infusion prior to open field tests, no catheter surgery was performed prior to intracranial surgery. Briefly, rats were mounted in a stereotaxic device. After making an incision in the skin on the skull, the nose bar was adjusted such that the change in dorsal-ventral coordinate from Lambda to Bregma was >0.1mm. We implanted the cannula at the following stereotaxic coordinates relative to Bregma: antero-posterior (AP) +1.3 mm; medio-lateral (ML) ±2.4 mm; dorsal-ventral (DV) -6.3 mm, 6° angle [5]. We anchored the cannula to the skull with 1/8" pan head sheet metal screws (Fastenere #842176107226) and dental cement (Stoelting #51458) and inserted blockers (Plastics One #C317DC) into the cannula barrel for protection.

*Intracranial virus infusion and fiber optic cannula implantation:* For rats destined for fiber photometry experiments, immediately following intravenous catheterization we performed bilateral intracranial infusions of virus expressing either the GRAB_DA2m biosensor (AAV9-hSyn-GRAB_DA2m, Addgene #140553) or the GRAB_DAmut control virus (AAV9-hSyn-GRAB_DAmut, Addgene #140555) into the NAcc followed by implantation of fiber optic cannula (Thor Labs, CFM15L10) bilaterally above each infusion site. Viruses were used after diluting 1:3 in sterile saline. Briefly, rats were mounted in a stereotaxic device and, after making an incision, the nose bar was adjusted such that the change in dorsal-ventral coordinates from Lambda to Bregma was <0.1 mm. In each hemisphere, we performed two infusions (250 nL each) of virus using the following stereotaxic coordinates (relative to Bregma): AP +1.3; ML ±2.4 mm; DV -7.2 mm for first infusion and DV -7.0 mm for the second infusion, 6° angle. We infused virus at 100nL/min and left the injector in place for 5 min following each infusion. Once the two infusions were completed, we raised the needle by 0.1 mm and waited 2 additional min prior to removal. Following the virus infusion, we implanted fiber optic cannula (AP +1.3 mm; ML ±2.4 mm; DV -6.8 mm; 6° angle). We anchored the cannula to the skull with 1/8" pan head sheet metal screws (Fastenere #842176107226) and dental cement (Stoelting #51458) and covered the fiber optic cannula with dust caps (Thor Labs, CAPF) for protection.

*Post-operative Procedures:* Rats received 5 mg/kg s.c. meloxicam (Covetrus, 6451602845, SKU #49755) as a post-operative analgesic. Rats for photometry experiments recovered for 4 days prior to beginning habituation/7 days prior to beginning SA (see Fig. 1A for timeline), and rats for pharmacology experiments recovered for 7 days prior to beginning SA (timelines in Figs. 4 and 6). For rats used for drug SA, catheters were flushed daily with cefazolin (0.1-0.3 mL of 0.1 g/mL in sterile 0.9% saline; Covetrus #54847) to prevent infection and maintain patency.

**Behavioral Apparatus**

*Self-administration chambers:* We trained and tested rats in Med-Associates (Fairfax, VT) self-administration (SA) chambers (#ENV-008-VPX) enclosed in sound attenuating chambers. Each chamber was equipped with two nose-poke holes (#ENV-114BM) on opposite sides of the apparatus which served as the operant manipulanda. The holes on the right side and left sides of the chamber represented the active and inactive ports, respectively. Photobeams (ENV-253SD) located at 1/3 and 2/3 of the length of the chamber were used to track activity. We used a single speed syringe pump (#PHM-100 or #PHM-108) that was mounted to the wall of the sound attenuating chamber or placed on top of the SA chamber to deliver infusions of intravenous cocaine. The pump was used to drive a 10 mL syringe that was connected (via a liquid swivel) to the rat's catheter using polyethylene-50 tubing protected by a metal spring.

*Open field:* Open field locomotion experiments took place in a 100 cm x 100 cm X 17 cm plexiglass open field (Stoelting 60200). We kept the apparatus 2 ft minimum from any other surface and elevated ~3ft above the ground to prevent escape from the apparatus.

### Behavioral Procedures

*Open field studies following intra-NAcc injection of DA receptor antagonists:* Open field studies were performed after recovery from surgery to implant bilateral intracranial guide cannulas. They began with 3 days of habituation to control for the effect of novelty on locomotion and accustom rats to the microinjection procedure. On the first day of habituation, rats were moved to the room where the microinjections would occur, placed on a desk for 30 sec of gentle head restraint, and then placed in their home cage for 15 min prior to being placed into the open field for 1 h. On the second habituation day, rats were returned to the room, their cannula blockers were removed, and then blockers were reinserted after rinsing sequentially in saline, 100% ethanol, and gentamicin (5 mg/mL diluted from 100mg/mL Covetrus, #6913). Rats again waited 15 min before being placed into the open field for 1 h. One the last day of habituation, the cannula blockers were removed and the injectors (Plastics One C317I) were lowered into the guide cannula and kept in place for 30 sec. Caps and injectors were rinsed as described above before reinsertion. Rats waited 15 min before being placed into the open field for 1 h. Rats received a day off prior to the first microinjection day and a day off in between microinjections (washout day) if two counterbalanced microinjections were given. On microinjection days, cannula blockers were removed. Injectors filled with drug or vehicle solution (an air bubble marked the start of the injected solution) were connected via Polyethylene Cannula Tubing PE50 10' (Plastics One, C313CT) to 10 µL Hamilton syringes (Hamilton, 80065) mounted in a syringe pump (World Instruments, UMP3 and Micro4). Injectors were inserted, extending 1 mm past the guide cannula into the NAc. Bilateral injection of vehicle or DA antagonist (1 µg/0.5 µL per hemisphere for both SCH39166 and L-741,626) was performed over 1 min. Injection was confirmed via air bubble movement in the tubing. Injectors were left in place for an additional min to allow for diffusion. Injectors were then removed and cannula blockers were reinserted after rinsing as described for habituation days. After 15 min, rats were placed in the open field apparatus and allowed to explore for 1 h. We recorded their behavior via a ceiling mounted webcam (Logitech C615) and OBS recording software (Version 27.2.3). Values for total distance traveled were generated with ANY-maze (version 7.16) video tracking software with a custom protocol. These values were then inputted into GraphPad Prism (version 10.2.2) and analyzed with an unpaired (SCH39166) or paired (L-741,626) t-test.

*Cocaine self-administration:* We trained rats to self-administer cocaine using an extended access regimen (6 h/day for 10 sessions; 5 days on, 2 days off, and 5 days on). The start of the session was indicated by the onset of white noise and a single cocaine infusion paired with port illumination. Following this, rats nose-poked in the active port to receive an infusion of cocaine (0.5 mg/kg/infusion) paired with 4 s port illumination which also represented the time-out period during which any additional nose-pokes were recorded but did not result in an infusion. We kept the concentration of cocaine in the syringe constant and varied the timing of infusion to correct for body weight (1.2-3.5 s) to deliver the same dose of cocaine to each rat. Responses in the inactive port were recorded but had no consequence. During SA training, food and water were provided ad libitum in the operant chamber. We tested rats that either failed to acquire SA (average infusions on SA days 8-10 < 25) or did not show greater responses in the active versus inactive port for catheter patency with 0.1 mL sodium brevital (Covetrus, #72465). Animals that failed the patency test were removed from the study (n = 2).

For virus group comparisons in the photometry experiments, we analyzed all data in GraphPad Prism (version 10.2.2) with mixed effects analysis with an assumption of sphericity (equal variability of differences). For assessment of nose poke behavior and discrimination between ports, we set fixed effects of virus (GRAB_DA2m or GRAB_DAmut), session (SA1-SA10), and port (active or inactive), with a random effect of subject. The factor of virus was between-subject while session/port were within-subject. To assess cocaine infusions, we performed another mixed effect analysis as previously described except with fixed effects of virus and session.

For analysis of any sex effects in photometry experiments we collapsed across virus groups and performed another mixed effects analysis, using same parameters as above, with fixed effects of sex (male or female), session (SA1-SA10) and port (active or inactive). The factor of sex was between-subject while session/port were within-subject. We assessed cocaine infusions via mixed effect analysis as previously described except with fixed effects of sex and session.

Fixed effects input into the model and output are specified in Table S2 and S3. When applicable we removed terms to fit a simpler model to our data. If multiple comparisons were run, we performed a Holm-Šídák correction on post-hoc analyses. Family-wise alpha threshold for confidence was set to 0.05.

*Cue-induced seeking tests during forced abstinence:* All rats received a cue-induced cocaine seeking test on forced abstinence day (FAD) 1 or 2, i.e., 24-48 h following the last day of SA (session 10). The experimental conditions during the seeking test and SA training were kept as similar as possible, except that active port responses now delivered the cue but no cocaine infusion. In addition, during the test rats did not have access to food or water and in photometry experiments rats were not tethered via their back port as in SA training. The number of responses on the active and inactive ports were recorded and active responses served as our measure of cocaine seeking. Locomotion during the test was monitored as photobeam breaks (see Apparatus). Following the FAD1-2 seeking test, rats underwent forced abstinence for 40-50 days in home cages (singly housed, as during SA). During forced abstinence, they were handled and weighed at least once a week, and estrous cycle monitoring was performed during certain periods as described below. Rats then underwent a second cocaine seeking test identical to the test performed on FAD1-2. For rats implanted with fiber optic cannulas, photometry measures were made throughout a 30 min test as detailed below under **Fiber photometry recordings.** Rats that received intra-cranial injections prior to testing had 1-h seeking tests. Data on the early and late FAD tests were compared via mixed effects analysis performed in GraphPad Prism (version 10.2.2) with an assumption of sphericity (equal variability of differences). For testing for an effect of virus expression on cocaine seeking, we set fixed effects of virus (GRAB_DA2m or GRAB_DAmut), test day (FAD1-2 or FAD40-50), and port (active or inactive), with a random effect of subject. The factor of virus was between-subject and test day/port were within-subject.

For analysis of any sex effects in photometry experiments we collapsed across virus groups and performed another mixed effects analysis, same parameters as above, with fixed effects of sex (male or female), test day (FAD1-2 or FAD40-50), and port (active or inactive). The factor of sex was between-subject and test day/port were within-subject.

Fixed effects input into the model and output are specified in Tables S2 and S3. When applicable we removed terms to fit a simpler model to our data. If multiple comparisons were run, we performed a Holm-Šídák correction on post-hoc analyses. Family-wise alpha threshold for confidence was set to 0.05.

*Cue-induced seeking tests preceded by intra-NAcc microinjections:* Rats for pharmacology studies received intra-NAcc infusions of DA receptor antagonists 15 min prior to either the FAD1 or FAD40-50 seeking test. These rats were habituated to the microinjection procedure as described above (*Open field studies following intra-NAcc injection of DA receptor antagonists*) except that rats were injected in a room separate to testing and then moved after the 15 min wait. On the day of the seeking test, bilateral infusion of vehicle or DA receptor antagonists into NAcc was performed 15 min prior to the start of a 1-h cue-induced seeking test. Rats were sorted into vehicle or treatment groups based on average infusions for the last 3 days of SA and, if being tested on FAD40-50, FAD1 active nose pokes. These animals were given a longer 1-h test to mirror similar studies from our lab [4] pokes on FAD40-50 were analyzed with a mixed effects model with a fixed effects of treatment (DA antagonist vs. vehicle) and port (active vs. inactive). Post hoc analysis of active port responding (DA antagonist vs. vehicle) was performed and corrected using Holm-Šídák tests.

We also analyzed data disaggregated for sex and ran the same analysis to examine possible sex differences.

**Fiber photometry recordings**

*Habituation:* For fiber photometry experiments, we began by habituating rats to the fiber optic cable 3 days prior to the start of SA training. We weighed each rat, cleaned their cannula with electronic grade 99.9% anhydrous isopropyl alcohol (MG Chemicals, #824-1L), and then placed the rat into the operant chamber. They were connected to a fiber optic patch cable (Thor labs Custom: FP-400URT, 0.5NA, 0.4m length, FT023SS tubing with a 2.5mm stainless steel ferrule) via a ceramic connector (Thor Labs, ADAF1-5) and the cable was passed through the top of the box and attached to a steel arm with a counterweight (Med Associates, PHM-110-SAI). The rats were then allowed to explore the operant box while tethered for 20 min after which they were returned to their home cage. Rats underwent 2 more days of habituation prior to SA. On the last day of habituation rats received highly palatable food pellets (LabDiet 5TUL) in the home cage to reduce neophobia when the same pellets were presented later in the experiment.

*Recording parameters:* Prior to all recordings, fiber optic cables were bleached at 200 mA for 8 h overnight. On the morning of the recording day, the LED power level and DC were adjusted so that both the 488 nm LED and 405 nm LED had a power output of 40 µW at the end of the fiber optic patch cable as measured using a Thor labs power meter (PM100D+). If the recording day lasted greater than 5 h, power output was checked for stability and changes were made to maintain 40 µW output. For recordings, we passed excitation wavelengths (488 nm and 405 nm) from the TDT RZ10x system via TDT patch cables (200 µm core, 2 m length, 0.5 NA). These TDT cables then connected to a Doric Minicube (FMC6) which in turn was coupled to two Thor fiber optic patch cables (FP400URT-Custom: FP400URT, FT0.5SS tubing, 4 m length coupled to a smaller 0.4 m patch cable via a TDT cable coupler) and then connected to the rat as described in *Habituation.* Emissions (500-550 nm and 460-490 nm) were received via the same patch cable and then decoupled at the Minicube and returned to the RZ10x via TDT response cables (600 µm, 2 m length, 0.5 NA). We acquired these emissions via the RZ10x photodetectors, digitized at 6kHz, and recorded in the TDT Synapse Software (version 95-44132P) on WS4 at frequency 330 Hz for 465 nm and 210 Hz for 405 nm. All recordings had a 6 Hz lowpass filter and a 9.5 V clip threshold.

*Prescreening:* Prescreening was conducted to verify that a DA signal was detected and identify the hemisphere with the best signal prior to advancing a rat to cue-induced seeking tests. We prescreened rats for DA signal on one of the two days off after SA session 5 and before SA session 6. For the prescreen, we cleaned the fiber optic cannula and then connected the rat to the recording set-up as described in *Recording parameters.* After 3 s of recording, the rat was placed in a novel context (Rubbermaid 71 qt container with same bedding as home cage) and allowed to explore for 1-2 min. Then, the rat was presented with highly palatable food pellets (Lab Diet 5TUL) and allowed to investigate and consume for 1 min. The rat was then removed from the novel context and the recording was repeated for the second hemisphere.

*Recording during cocaine seeking:* For recordings during cue-induced seeking tests, the rat was removed from the home cage, weighed, and had their fiber optic cannula cleaned. We connected the fiber optic patch cable via a ceramic connector as described in *Recording parameters*. The patch cable was passed through the SA chamber on a counterweight system as described in *Habituation*. Rats were placed into the operant chamber and the photometry recording began. The Med Associates program (to record nose-pokes and infusions) was started 5 s later to avoid light artifact obscuring any start of session signals. The photometry recording was continuous for the entire 30-min seeking test. During the test, rats were also recorded via a Logitech webcam (#C615) and checked every 10 min for possible disconnection or tangling of the fiber optic cable. If an intervention was required, it was done as quickly and unobtrusively as possible and the time of the intervention during the test session was noted. After the seeking test ended, the photometry recording was halted and the rat was returned to the home cage.

*Data analysis:* Fiber photometry data were analyzed using the analysis suite GuPPy created by the Lerner Lab and can be downloaded on their GitHub page; the details of the data manipulations used to generate the GRAB_DA2m signal traces are described in [6]. In brief, for each recording session, we applied least-squares linear fit to align isosbestic and signal channels to one another and use fluctuations in the isosbestic channel to account for florescence changes in the signal channel that are not a result of DA binding. Any disconnects during the recording, defined by a rapid substantial shift (>10 mV) in average mV, were snipped prior to fitting the traces and any nose pokes within those snips were excluded from further analysis. Then we calculated a change in fluorescence measure (ΔF/F=Signal-Fitted Control/Fitted Control) and applied z-score normalization to control for between-session and between-rat differences in virus expression or recording. Finally, we averaged normalized traces across behavioral event replicates for each rat, then averaged by day (early or late seeking test).

For analysis of the entire session trace (Fig. 2E), we examined the average spike amplitude (spikes are defined as transients that are >7 median absolute deviations above the median of the 15-s moving window, regardless of whether they coincide with a nose-poke) as well as the frequency of these spikes during the seeking tests. This threshold was determined by analyzing our GRAB_DAmut recordings and selecting parameters that lead to zero spike detection in these recordings. We analyzed these whole session values with a mixed effects model to assess whether test day had a significant effect on frequency or average spike amplitude.

For DA transients associated with nose-pokes, first we extracted z-scored ΔF/F traces within a 6-sec window around each behavioral response (active or inactive nose-poke) and averaged all traces to generate an average response for the rat during each test. We identified significant DA transients using continuous threshold bootstrapping methods. Bootstrapping is a statistical method in which multiple simulated data sets are generated from an original data set to enable hypothesis testing and calculation of confidence intervals. When applied to fiber photometry data we consider each timepoint in the trace to be a data set, with each rat contributing 1 data point from their individual averaged trace. We then generate 1000 replications of this data set by resampling with replacement from our original 17 data points (one for each rat). This generates a normal distribution of values for each timepoint, which can then be used to generate confidence intervals. These confidence intervals are then used as a method to detect likely transients (i.e., 95% likely z-score ≠ 0) in the entire 6-sec time window analyzed. However, rapid non-physiological changes in fluorescence can occur, for example, due to a cable hitting a wall. To account for these non-physiological events, we set a consecutive threshold where the 95% confidence interval cannot contain 0 (baseline) for a consecutive number of samples set in the code parameters. We selected a consecutive threshold of 125 for our 6 kHz acquisition, which equates to 0.02083 s. This was based on applying this analysis to recordings from rats expressing GRAB_DAmut and selecting a threshold which resulted in no significant transients. Bootstrapping is thought to provide an unbiased way to identify relevant transients in photometry without experimenter selection of a time window and can be considered a data driven approach. In tandem with this analysis, we also performed the classical hypothesis driven approach in which we selected time windows before and after our behavioral event of interest and calculated the Area Under the Curve (AUC) for the z-scored ΔF/F trace within this time window. Using these AUC values, we performed a linear mixed effects model with factors of test day (FAD1, FAD40+) and sex (M, F). We analyzed DA responses separately for active pokes that triggered the cue, active pokes during the 4-s cue/time-out period, and inactive pokes. We also analyzed the data after separating the 30-min test into three 10-min bins to assess potential within-test changes in DA responding.

#### Estrous cycle monitoring

We performed estrous cycle monitoring as but females in the estrus phase show significantly increased seeking compared to makes and non-estrus females [7-9]. This potentiated seeking could be due to the sex hormone estradiol modulating DA-R signaling [10]. Additionally, female mice in estrus show increased phasic activity of VTA DA neurons measured using single-unit recordings [11]. We can't rule out small differences in DA release, not detectable with our methods, related to the estrous cycle.

Therefore, Estrous cycle monitoring was conducted for 4-8 days around both the FAD1-2 and FAD40-50 seeking tests. The monitoring consisted of weighing the animal and, if female, performing a vaginal lavage; if male, we gently prodded the testes as a control manipulation. To habituate rats to these procedures, they were also performed (but estrous cycle stage was not scored) once a week during the forced abstinence phase. Vaginal lavages were performed as described [12]. Briefly, female rats were weighed and then held with their tail lifted to expose their genitalia. A glass blunt-tipped eye dropper (TecUnite, 1mL) containing ~0.3-0.5 mL sterile 0.9% saline was placed at the entrance of the vagina; the saline was quickly inserted and withdrawn. This saline was then placed in a labeled 24-well plate and kept at 4˚C for a maximum of 4 days prior to image analysis. Lavages were imaged in the 24-well plate. Images were acquired at 20 X magnification using a Leica DMi8 inverted microscope equipped with an ORCA-Flash4.0 LT+ Digital CMOS camera (Hamamatsu). Leica Application Suite X (LASX) Premium Software (version 3.7.5.24914) was used for image acquisition and ImageJ was used for analysis. After all images were acquired, we examined the lavages for each rats across days and assigned the lavage as estrus (identified by large numbers of cornified cells that lack nuclei), proestrus (identified by large numbers of nucleated cells), diestrus (identified by large numbers of lymphocytes), or metestrus (identified by a mix of lymphocytes and cornified cells) [12]. It should be noted that incubation studies show estrus effects used between-subject designs. A study using a within-subject design, similar to our study, found no differences in incubation of cocaine craving between males and females or between females in estrus and non-estrus phases of the cycle [13].

#### Histology

Upon completion of experiments, rats were euthanized via lethal injection of Fatal Plus (Covetrus, #35946) diluted to 80 mg/kg with sterile saline. Once animals no longer displayed reflexive motor responses, they were perfused transcardially first with 1x phosphate buffer saline (PBS) followed by 4% formaldehyde/1% methanol in 1x PBS, pH ~7, at a rate of ~100 mL/s over 5 min. Brains were extracted and allowed to rest in the formaldehyde solution for up to 24 h. Brains were transferred to 1x PBS with 0.01% sodium azide and then sliced on a vibratome (Leica VT1000s; frequency 8, speed 70, blade DORCO plat. 5T300) at 60 µm. Slices were kept in a 24 well plate in 1x PBS with 0.01% sodium azide at 4˚C until the start of immunohistochemistry or cresyl violet staining.

*Cresyl violet staining:* Brains from intracranial DA receptor antagonist microinfusion experiments were mounted on Superfrost Plus slides (Fisher Scientific, Cat# 1255015) after slicing and allowed to dry completely. Once dry the tissue was rinsed on the slides in decreasing concentrations of EtOH (100%, 95%, 70%, 0%) for 3 min in each concentration. The slides were then placed in a Cresyl violet acetate solution (1 g Cresyl Violet Acetate/2.5 mL 100% Glacial Acetic Acid/1 L $H_2O$) and remained there for 2-4 min or until the stain was sufficiently dark. After staining, slides were placed into increasing concentrations of EtOH (0%, 70%, 95%, 100%) for 15 sec each. Next the tissue was rinsed with CitriSolv Hybrid (Decon Labs Inc., Cat #1601H) twice for 5 min each. Tissue was coverslipped (Fisher Scientific, 12541026) directly after the last CitriSolv rinse with Permount (Fisher Scientific, Cat #SP15-100) and allowed to dry for 2 days.

*Immunohistochemistry:* We performed immunohistochemistry (IHC) on tissue from photometry experiments to amplify the GFP signal prior to imaging to confirm virus expression and placement. We began IHC with three 30-min washes, first in 1x PBS (diluted from 10x PBS, Quality Biological, 119-069-151) then twice in 1x PBS with 0.5% (v/v) Triton-X100 (Electron Microscopy Sciences, #22140). After this we permeabilized the tissue for 2 h in 1x PBS with 0.5% (v/v) Triton-X100, 20% (v/v) DMSO (Sigma Aldrich, #276855), and 2% (w/v) Glycine (Sigma Aldrich, #G8898) at room temperature (RT) on a rocking shaker. Next, we blocked tissue in 1x PBS with 0.5% (v/v) Triton-X100, 10% (v/v) DMSO, and 6% (v/v) Normal Donkey Serum (NDS, Jackson Immuno Research, 017-200-121) for 2 h at RT on a rocking shaker. After blocking we incubated the tissue overnight at RT in 1:1000 Anti-GFP (Aves, GFP-1010) in 1x PBS with 0.5% (v/v) Tween-20 and 0.01% (w/v) Heparin (Sigma Aldrich, #H3393-100KU) with 3% NDS and 10% DMSO on a rotator. The following day, we washed the tissue 3 times for 30 min each in 1x PBS with 0.5% (v/v) Tween-20 (Thermo Scientific, #J20605-AP) and 0.01% (w/v) Heparin before incubating in 1:250 Anti-chicken-488 (Jackson Immuno Research, 775-546-155) in 1x PBS with 0.5% (v/v) Tween-20 and 0.01% (w/v) Heparin with 3% NDS overnight at RT on a rotator. The third day, we performed two 30-min washes in 1x PBS with 0.5% (v/v) Tween-20 and 0.01% (w/v) Heparin before beginning a final wash in 1x PBS (30 min). Slices were kept in 1x PBS with 0.01% sodium azide at 4˚C until being mounted onto Superfrost Plus slides and coverslipped using Vectashield Vibrance with DAPI (Vector Labs, H-1800-10).

*Imaging* Images were acquired using a Leica DMi8 inverted microscope equipped with an ORCA-Flash4.0 LT+ Digital CMOS camera (Hamamatsu). LASX Premium Software was used for image acquisition and ImageJ was used for analysis. Exposure was determined using the software to avoid saturation while maximizing the pixel distribution in the histogram. For DAPI and FITC at 2.5X, we used a 3-s exposure.
